# An expert-based global assessment of threats and conservation measures for subterranean ecosystems

**DOI:** 10.1101/2023.01.09.523190

**Authors:** Veronica Nanni, Elena Piano, Pedro Cardoso, Marco Isaia, Stefano Mammola

## Abstract

Subterranean ecosystems host unique biodiversity and deliver important services to humans. Yet, available data for subterranean ecosystems are limited in space and/or taxonomic scope and global monitoring programs are absent, preventing practitioners to develop effective conservation and management strategies. Expert opinion may help overcome some of these knowledge gaps. We designed a global survey to quantify the importance of anthropogenic impacts and conservation measures for subterranean ecosystems. We obtained 279 responses from 155 experts in different subterranean habitats, taxa, and regions. Experts perceived surface habitat change, direct habitat destruction (e.g., pollution, damming, mining), and climate change as the most relevant threats impacting subterranean ecosystems. Legislation, land protection, and education were scored as the most effective conservation measures, whereas species-level conservation was deemed less relevant. Whenever lacking hard data, expert opinion may be an effective, largely available, yet often overlooked source of information to implement timely conservation interventions for subterranean ecosystems.

## INTRODUCTION

Subterranean ecosystems—including caves, fissure systems, and groundwaters—harbor a broad diversity of poorly-understood specialized organisms that are of interest from both an evolutionary and conservation perspective (Culver & Pipan, 2019). Subterranean-dwelling species are often phylogenetically highly distinct, yet small-range endemics, and some are representative of ancient faunas that have disappeared from above surface habitats. Thus, they account for a unique fraction of the global taxonomic, phylogenetic, and functional diversity (Mammola, et al., 2019b). Furthermore, subterranean ecosystems and landscapes deliver important ecosystem services (Canedoli et al., 2022), including provisioning (e.g., drinking water, guano), regulating (e.g. water quality, shelter for predator taxa), supporting (e.g., soil formation) and cultural (recreational caving, tourism, education) ones. An estimated 95% of the world’s available liquid freshwater supply is groundwater, and more than half of the world’s population relies on this supply for consumption and agriculture (Griebler & Avramov, 2015). Subterranean bacteria and invertebrates are essential in maintaining clean groundwaters by enhancing carbon turnover, attenuating and degrading harmful contaminants, and even eliminating pathogenic microorganisms from the aquifer (Boulton et al., 2008; Herman et al., 2001). But the services provided by subterranean organisms extend beyond that. For example, bats form the largest congregations of mammals roosting in caves, playing crucial ecological roles as insect pest controllers, seed dispersers, pollinators, and nutrient recyclers (Kunz et al., 2011). This well-illustrates the importance of subterranean ecosystems to the integrity of ecological systems (including surface ones) and to human societies, but also reminds us about how little we still know about these secluded environments (Ficetola et al., 2019).

Anthropogenic pressure on subterranean ecosystems is escalating, with multiple threats affecting species and habitats at different spatial-temporal scales and often with either cumulative or synergistic effects (Mammola, Cardoso, et al., 2019; Raghavan et al., 2021; Wynne et al., 2021) The joint effect of surface land use change and climate change represents a major impact to subterranean ecosystems at different scales. Indeed, modification of habitats at the surface (e.g., urbanization, agricultural activities, deforestation) may feedback to affect subterranean ecological dynamics and climatic conditions. For example, loss of surface vegetation due to deforestation and wildfires can quickly lead to broad-scale habitat alterations (e.g., desertification), changing water balance and altering temperature of subterranean atmospheres (Sánchez-Fernández et al., 2021; Shu et al., 2013; Whitten, 2009). These impacts are further exacerbated by global warming, directly modifying underground climates (Badino, 2004), reducing groundwater availability (Wu et al., 2020), and impacting specialized species with reduced thermal tolerance and low dispersal capacity (Mammola, et al., 2019). Additionally, at local scales, countless threats have been documented to potentially affect specific subterranean habitats and/or species, including habitat loss (e.g., mining activities; Ferreira et al., 2022; damming; Fišer et al., 2022), local polluting events (Manenti et al., 2021), diseases (Hoyt et al., 2021), poaching (Simičević, 2017), alien species introductions (Nicolosi et al., 2023), and even stochastic events (e.g., floods; Pacioglu et al., 2019; earthquakes; Fattorini et al., 2018). This cocktail of threats, coupled with the intrinsic vulnerability of subterranean species due to their ecological specialization, makes the protection of subterranean ecosystems a challenging endeavor.

Recently, we reviewed available knowledge on the effectiveness of subterranean conservation interventions considering a breadth of threats, organisms, and systems (Mammola et al., 2022). From this synthesis, it emerged that the evidence-base for subterranean conservation is far from being organized and exhaustive. Although some conservation efforts have been devoted to protecting specific species or habitats at local scales (e.g., Manenti et al., 2019; Turner et al., 2022), no global assessment addressing the challenge of preserving and protecting subterranean biodiversity exists so far. Large-scale studies and long historical series of data are scarce (e.g., Tanalgo et al., 2022), and our knowledge is biased in its taxonomic and geographical coverage (Mammola et al., 2022). Ultimately, this lack of quantitative understanding of population trends for subterranean species and the threats affecting them is a central impediment to conservation.

As global monitoring programs and unbiased data are poorly available, we must explore alternative approaches to advance our knowledge on subterranean ecosystems. In the case of other data-poor scenarios, expert opinion emerged as a valuable tool for consolidating a first understanding on current threats and conservation needs (Branco & Cardoso, 2020; Chamberlain et al., 2016; Fitzgerald et al., 2021; Luo et al., 2023; Miličić et al., 2021; O’Neill et al., 2008). Indeed, when lacking hard data, expert opinion plays a fundamental role in conservation (Martin et al., 2012), e.g. by informing decision-making (Cook et al., 2010) and risk assessments (Patterson et al., 2007).

Here, we surveyed experts on a wide variety of subterranean ecosystems and taxa around the world, aiming to quantify the importance of anthropogenic threats and conservation measures for subterranean ecosystems and to provide a roadmap on how to preserve these fundamental habitats and associated species and ecosystem services.

## METHODS

### Expert survey

We created an online questionnaire in Google Forms and disseminated it to experts on subterranean biology globally (April–September 2021). A limitation in expert opinion surveys is that experts tend to be systematically overconfident, i.e. their subjective probability distributions tend to be too narrow (Granger Morgan et al., 2001). To minimize the bias so introduced, we attempted to maximize the academic age and degree of experience of the respondents while keeping the definition of what constitutes an expert broad. Specifically, we considered an “expert” any respondent knowledgeable of subterranean taxa or habitats, including research, applied science, decision making, and/or education. Furthermore, to maximize geographical coverage, we contacted global societies and journals focusing on subterranean biology (e.g., International Society for Subterranean Biology, Subterranean Ecology, Subterranean Biology Journal) and asked them to distribute the query by email among their members. To ensure the sample stratification among subterranean taxa, we also contacted local societies (e.g., African Bats Conservation, North American Society for Bat Research, BatLabFinland, Chirosphera, Società Erpetologica Italiana, Italian Spiders; Speleological Survey Group of Yamaguchi University, Biologia Sotterranea Piemonte, Speleovivarium Trieste, Laboratory of Subterranean Biology “Enrico Pezzoli”, Cave Conservation Australia, Institute of Speleology-Romanian Academy). Finally, we shared the questionnaire through social media (ResearchGate, LinkedIn, and Twitter). We estimated the questionnaire reached an upper boundary of 350–400 recipients.

We structured the questionnaire into three main sections (Supplementary Material 1). In the first section, questions 1− 3 aimed to provide information on the demographic structure of the respondents and their expertise (education, experience, and work responsibilities). Questions 4− 7 were intended to reveal their biogeographic, habitat, and taxon expertise. If the respondents’ expertise covered several biogeographic regions or taxa, we encouraged them to fill the survey multiple times, one for each combination. For the habitat, respondents could select multiple habitats expertise and we subsequently splitted the answers. We eliminated answers from respondents which selected “I don’t have any expertise on subterranean ecosystems” (n = 8). In the second section (question 8–12), respondents gave information regarding threats and conservation measures on their taxonomic group of expertise, ranking their importance according to the Linker scale from 1 (“Not relevant”) to 5 (“Most relevant”) or indicating the option “I don’t know” if uncertain. We selected threats for subterranean environments and conservation measures based on a recent systematic review on the conservation of subterranean ecosystems (Mammola et al., 2022), as well as similar expert-opinion surveys (Branco & Cardoso, 2020; Miličić et al., 2021). In the last question, we asked respondents to provide references supporting their answers and to leave any comments or suggestions.

Prior to dissemination, we piloted the survey within a selected group of colleagues specialized in subterranean biology, asking for feedback on the survey’ structure and on the clarity of questions. Indeed, a typical bias in expert surveys may occur because of unclear questions which may be understood differently by the respondents. As a result, we made minor changes to the phrasing of questions and the overall structure of the questionnaire.

### Statistical analyses

We ran all analyses in R version 4.1.0. We fitted regression models to predict the importance scores assigned to each threat and conservation measure while controlling for confounding factors related to respondents’ education and experience with subterranean biology, meanwhile testing for variations in responses among biogeographic regions and taxa. Since the dependent variable was an ordinal score (ordered factor with five levels from “Not relevant” to “Most relevant”), we fitted regressions for ordinal data within a bayesian framework using the R package ‘brms’ version 2.15.0 (Bürkner, 2017; Bürkner, 2018). The choice of a Bayesian model was driven by a general scarcity of R packages able to handle ordinal data with a frequentist approach.

We set a cumulative error family and a logit link function with an equidistant threshold among the ordered scores. Model parameters were as follows: 4 chains, 2000 iterations, 1000 iterations as burn-in, default priors, a maximum tree depth of 15, and an adapt_delta of 0.99— which is computationally slower but avoids divergent transitions after warmup. The structure of the 20 models (14 models for threats and 7 for conservation measures) was:

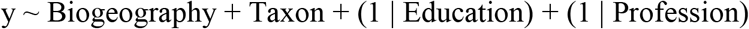

Whereby we tested for differences among biogeographic regions (Biogeography, a categorical variable with 6 levels; Figure 1A) and organisms (Taxon, 7 levels), and we included two random intercept factors that accounted for confounding effects of respondents’ level of education (Figure 1B) and professional status (Figure 1C). In other words, the random factors controlled for the fact that experts with the same education level and professional status were likely to provide more similar votes than expected from random. Note that we did not include academic age (Figure 1D) and habitat type (Figure 1A–D) in the model, being these variables significantly associated with Education and Taxon, respectively.

**Figure 1.**
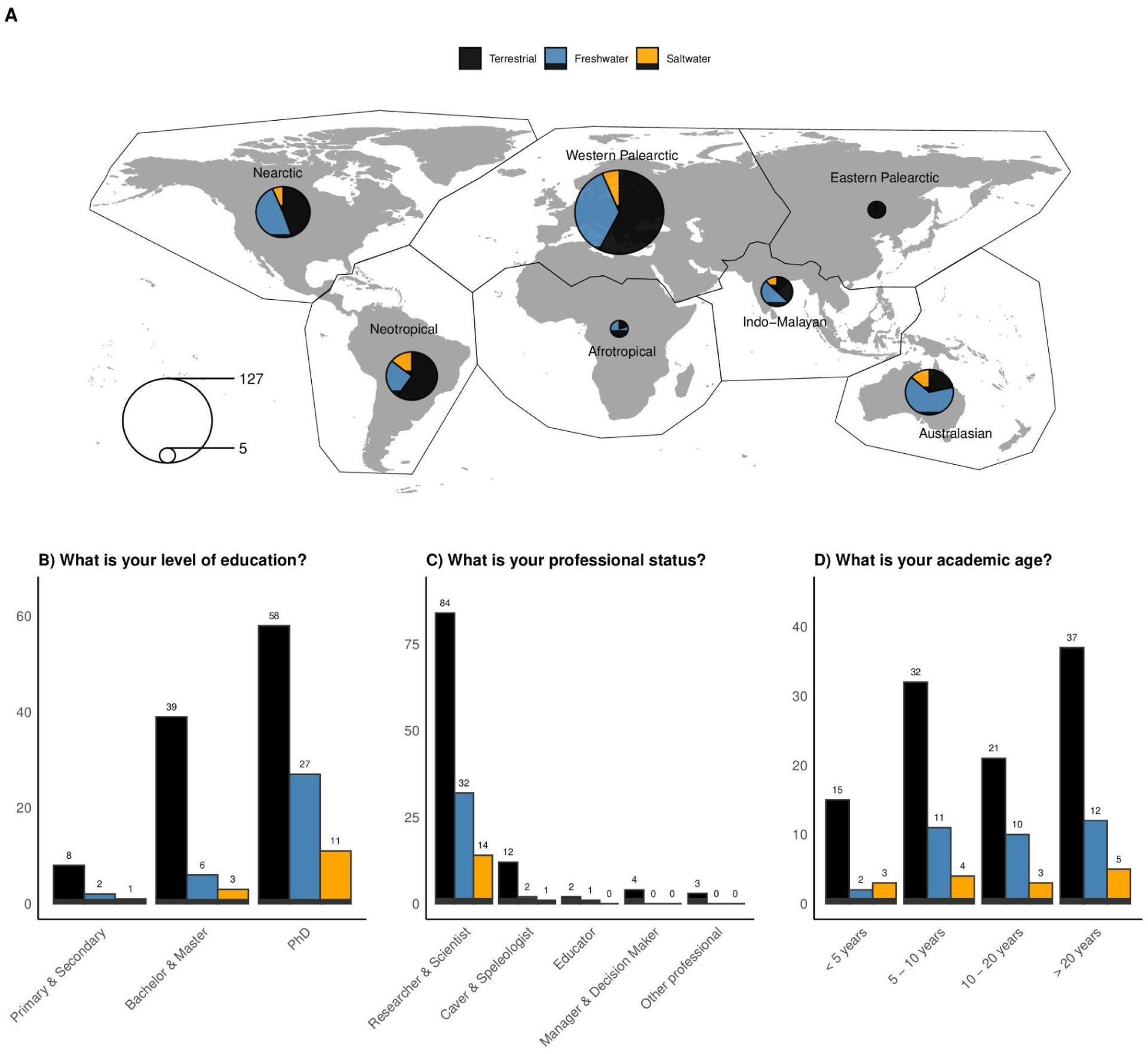
General overview of the survey. **a**) Number of responses by biogeographic regions; **b**–**d**) Breakdown of experts by level of education, professional status, and academic age.

We validated models by inspecting the mixing of chains—we detected no divergences except for the dependent variables “Other threats” and “Other conservation measures”, which had much reduced sample size. We used the final models to predict the votes for each individual respondent for the full dataset and also for the biogeographic regions and taxa, thereby obtaining scores corrected for the confounding factors above. Predictions for ordinal regression are probability values for each level of the ordinal dependent variable. See supplementary material for a comparison between predicted and observed values for the scores of threats (Figure S1) and conservation measures (Figure S2).

## RESULT & DISCUSSION

### The survey by numbers

We obtained a dataset of 279 responses from 155 participants. There was uneven geographic coverage, with most answers coming from the Western Palearctic (45.5%, n = 127) (Figure 1A). The best represented group of subterranean organisms were aquatic and terrestrial invertebrates (30.5%, n = 85 and 23.7%, n = 66, respectively). In agreement with our target audience, most respondents had a PhD (62.7%, n = 175) (Figure 1B) and were academics and researchers (86%, n = 234) with recreational cavers or speleologists (9.9%, n = 27), decision-makers (2.6%, n = 7), and educators (1.5%, n = 4) also being represented (Figure 1C). The level of education was generally high with only 6.8% of respondents not graduated; the majority of respondents had >20 years of expertise on subterranean ecosystems (35.8%, n = 100) (Figure 1D).

### Expert perception of threats

At the global level, threats perceived by experts as most relevant for subterranean ecosystems are related to surface (agriculture, forestry or urbanization) and subterranean habitat change (pollution, water management, and mining) (Figure 2A). According to experts, subterranean ecosystems are also highly threatened by climate change, transversely affecting geographic regions and taxa. Conversely, poaching, tourism, wildfires, and geological events were perceived as less important threats. Poaching and tourism are occasional events affecting single caves and species populations (Piano et al., 2022). Poaching in caves is poorly studied (Simičević, 2017), and hence expert opinion may be driven by poor awareness. Geological events are mostly unpredictable and probably have a tangible effect on subterranean fauna in a few cases, such as large volcanic eruptions on oceanic islands that have the potential to wipe entire species but remain relatively rare events. Furthermore, even if some natural catastrophic events are exacerbated by climate change and land use practices, there is very little conservation biologists can practically do to prevent them.

**Figure 2.**
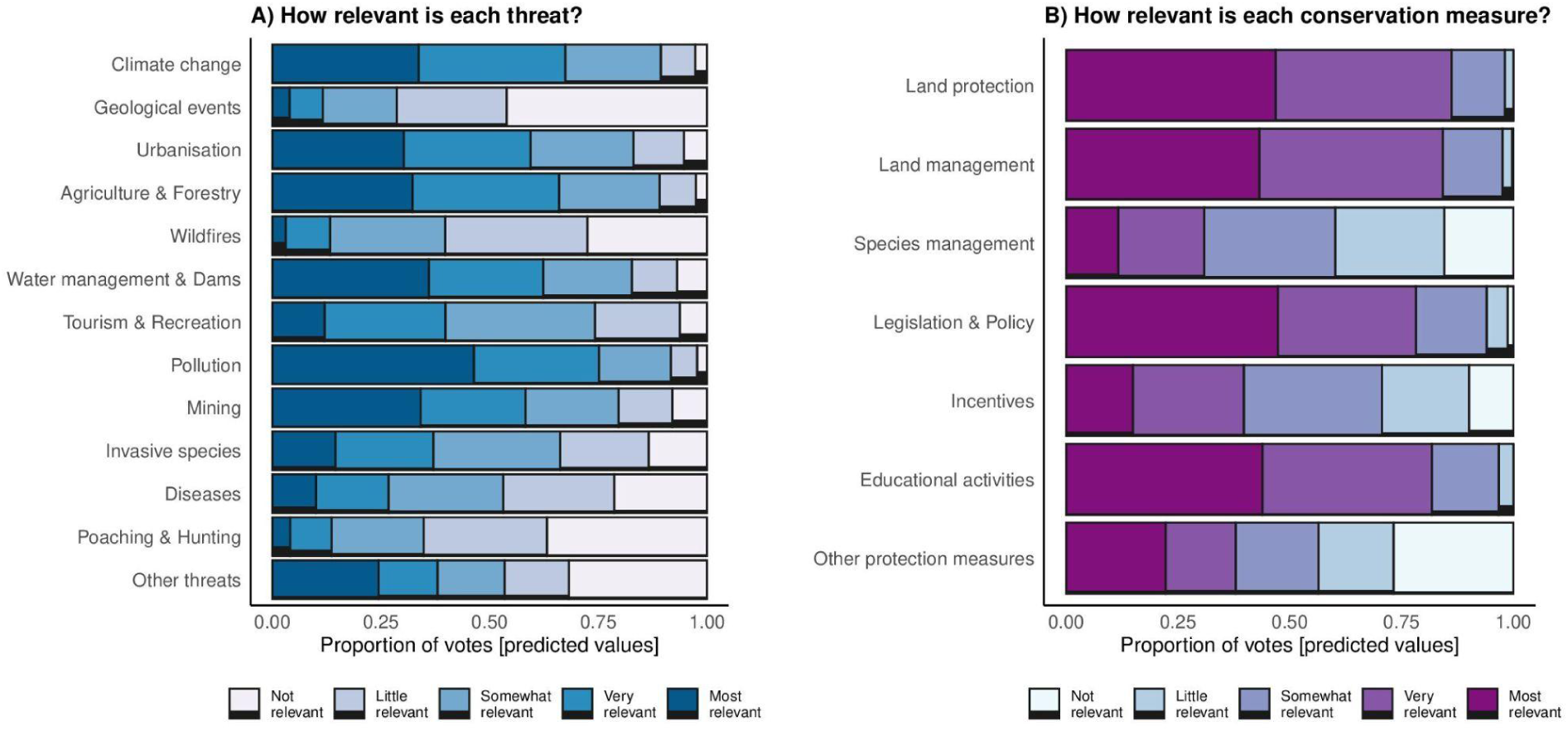
Importance of each threat (A) and conservation measure (B) based on the responses by surveyed experts. Proportion of votes is expressed according to a Linkert scale. Note that scores are not the original values, but predictions from regression models that control for confounding factors related to respondents’ education and degree of experience with subterranean biology. For each threat and conservation measure, Supplementary Figures S1–S2 reports the observed scores vs the scores predicted by the ordinal regression models.

Respondents also indicated other threats that may concur to determine the decline of subterranean habitats and taxa. A frequently mentioned problem was the government’s complacency and inaction towards the environment, which is often aggravated by widespread economic interests on the exploration of minerals often associated with karstic regions. Economic pressures are increasing especially in developing countries, and those are not properly counteracted by environmental laws—which sometimes do exist, but are rarely enforced when economic interests are prevalent. A recent problem mentioned by experts was the sanitation of tourist caves against COVID-19, which may be detrimental depending on chemicals and application methods (see Barton, 2020). Finally, some experts emphasized the importance of context-specific threats related to local traditions and culture—for example, the vulnerability due to religious activities in regions where caves are used as temples or praying sites.

Importantly, there was some variation in threats identified as the most relevant across taxa (Figure 3) and biogeographic regions (Figure 4). In general, terrestrial vertebrates were perceived as less sensitive to pollutants, climate change, and agriculture/forestry compared to other taxa (Figure 3). Bats differed from other taxa with respect to the impact of tourism, recreation and wildfires which, according to experts, affect them disproportionately more. Bats sensitivity to diseases mostly pertains to the white nose syndrome, a disease caused by the fungus *Pseudogymnoascus destructans* which has led to mass mortality of cave-roosting species across North America (Hoyt et al., 2021). Furthermore, bats’ sensibility to human disturbance in caves is well-documented (Tanalgo et al., 2022; Voigt & Kingston, 2016) and wildfires are becoming a pressing problem in the last decades aggravated by climate change (Flannigan et al., 2009). Microorganisms were perceived as particularly vulnerable to pollution, agriculture, and forestry. For example, nitrogen fertilizers used in agriculture may alter the microbial community and the biogeochemical cycling of nitrogen within aquifers (Marschner et al., 2003). Concerning geographic variations, pollution and water management were perceived as less important for Eastern Palearctic, while agriculture/forestry was scored higher for Indo-Malayan and Neotropical regions (Figure 4). Urbanization was not perceived as a major threat in Afrotropical and Australasian regions, where its pressure is generally lower.

**Figure 3.**
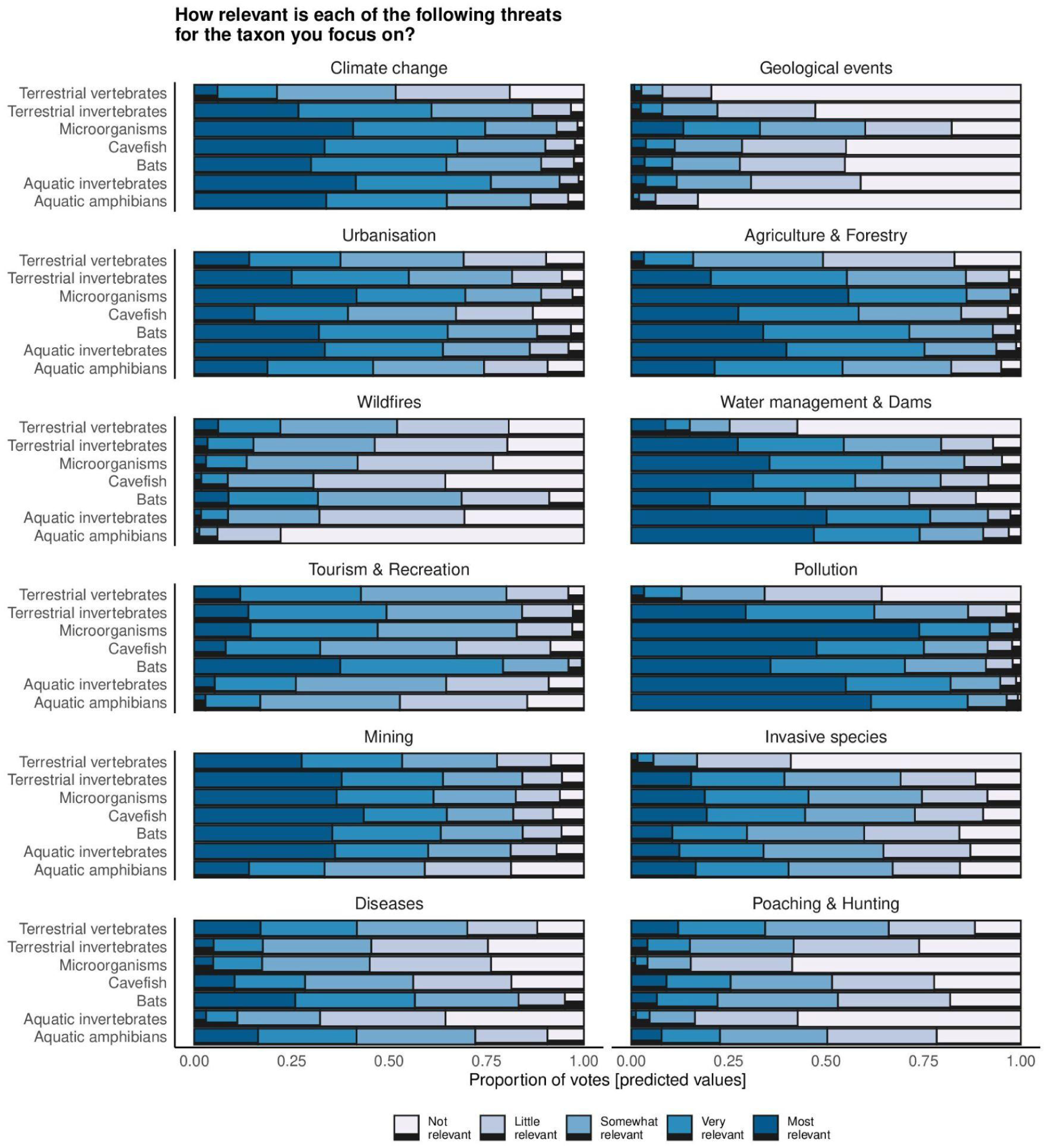
Importance of each threat across different taxa based on the responses by surveyed experts. Proportion of votes is expressed according to a Linkert scale. Note that scores are not the original values, but predictions from regression models that control for confounding factors related to respondents’ education and degree of experience with subterranean biology. Supplementary Figures S3–S21 show the direction of effects for each threat and conservation measure according to ordinal regression models, while model estimates are available in Supplementary Tables S1–S20.

**Figure 4.**
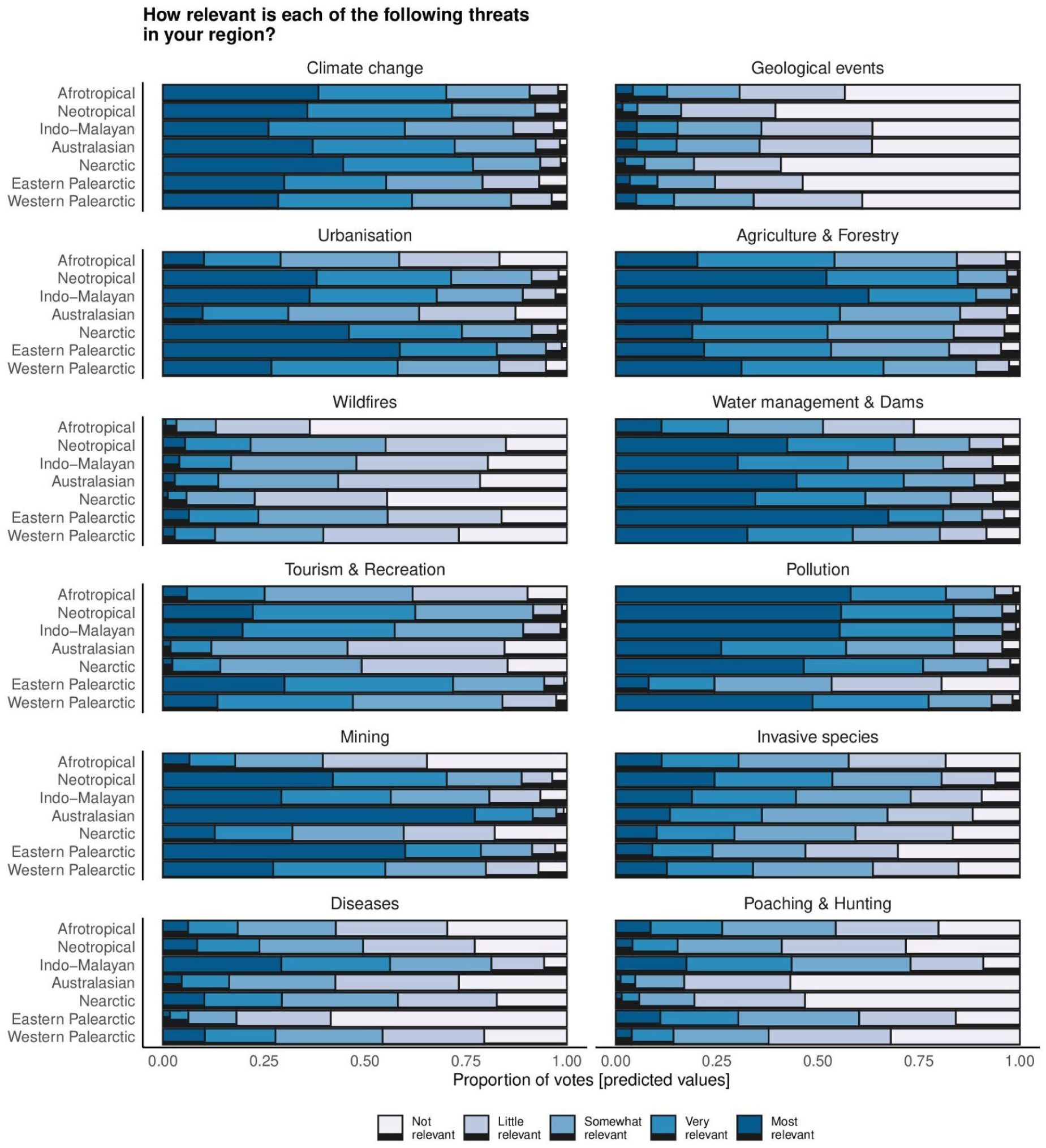
Importance of each threat across biogeographic regions based on the responses by surveyed experts. Proportion of votes is expressed according to a Linkert scale. Note that scores are not the original values, but predictions from regression models that control for confounding factors related to respondents’ education and degree of experience with subterranean biology. Supplementary Figures S3–S21 show the direction of effects for each threat and conservation measure according to ordinal regression models, while model estimates are available in Supplementary Tables S1–S20.

### Expert perception of conservation measures

Experts identified legislation and policy jointly with land protection and management as the most relevant conservation measures (Figure 2B). Area-based conservation (i.e., the establishment of legally protected areas and their correct management) is widely regarded among the most effective strategies for halting biodiversity loss across different ecosystems (Maxwell et al., 2020). Reserving large areas for nature is especially effective whenever there is a poor knowledge of the real extent of the habitat to be protected and the ecology of species therein, as is often the case for “out of sight” subterranean ecosystems (Ficetola et al., 2019; Mammola et al., 2019a). Currently, area-based protection of subterranean ecosystems is mostly indirect, namely when subterranean resources overlap with surface protected land such as national parks and nature world heritage areas (Sánchez-Fernández et al., 2021). However, according to recent estimations, the current network of protected areas largely fails to protect subterranean biodiversity (Colado et al., 2022), emphasizing the importance of establishing protected areas and legislations able to account for the 3-dimensionality and specificities of subterranean ecosystems (Iannella et al., 2021).

Experts also greatly valued the role of education in supporting long-lasting conservation outcomes. This result is a powerful reminder that conservation can only progress if humans value nature and care for it. Effective educational practices span indirect and direct actions, including place-based experiences in natural contexts, as well as building a community with shared environmental norms which actively take part in conservation actions (Niemiec et al., 2016). The involvement of local communities could also happen through new job opportunities, e.g. by engaging environmental rangers in karst areas protection, which will create economical benefits to the community itself. Ultimately, with education comes awareness and, in the long run, an empowerment of local communities to stand for local biodiversity and natural resources. This is largely perceived as the most effective way to ensure that environmental legislation is concretely implemented and respected (Ardoin et al., 2020).

Conversely, experts perceived species-level conservation and management as less relevant. This is probably linked to the fact that subterranean species often have narrow-range distribution, preventing the effective protection of several individual species or to select suitable widely distributed umbrella taxa whose protection would benefit a large number of species. Moreover, our knowledge on the distribution and state of conservation of underground organisms is scarce for species management planning. The cost-effectiveness of species-level conservation can also be low, as measures targeting individual species are often not effective for other species if the full ecosystem is not somehow protected or restored. Some measures such as ex-situ conservation, breeding and eventual repopulation can however be further explored for particular cases.

Once again, there was some variation in conservation actions identified as the most relevant across taxa (Figure 5) and biogeographic regions (Figure 6). Incentives were scored as an important conservation measure for terrestrial and aquatic invertebrates, microorganisms, cavefish, and bats. Given that these organisms are less charismatic than terrestrial vertebrates and amphibians, it may be that they are less targeted by financial investment into conservation. Education was scored as a less important activity in the Afrotropic and Eastern Palearctic. Land protection and management were deemed particularly important for the Afrotropic and Indo-Malaya, possibly because land modifications in these regions are occurring rapidly with lack of regulations and management. Finally, species management was suggested to be important for cavefish conservation, possibly reflecting the fact that harvest, trade, and poor groundwater governance are emerging concerns in different areas (Raghavan et al., 2021).

**Figure 5.**
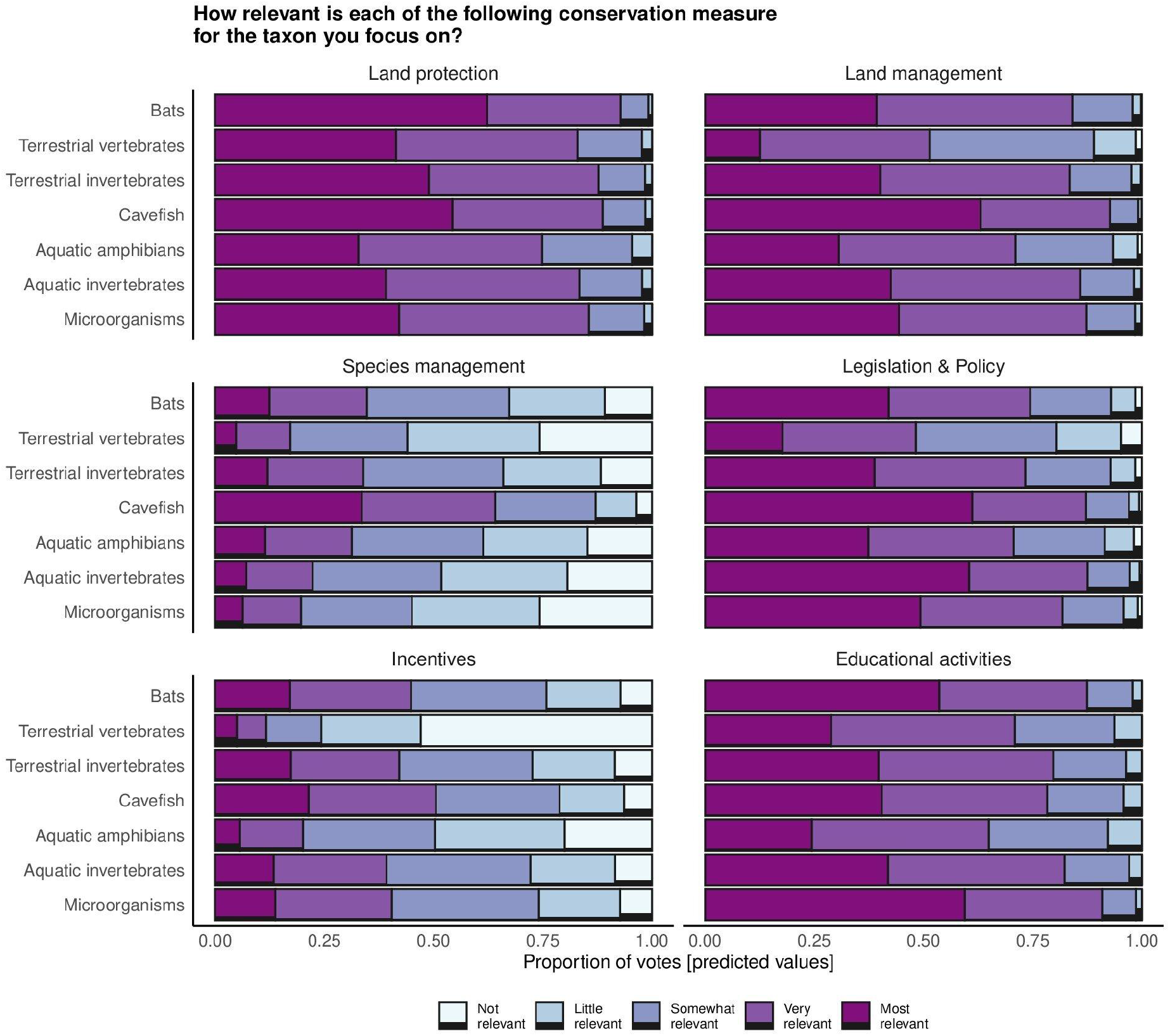
Importance of each conservation measure across different taxa based on the responses by surveyed experts. Proportion of votes is expressed according to a Linkert scale. Note that scores are not the original values, but predictions from regression models that control for confounding factors related to respondents’ education and degree of experience with subterranean biology. Supplementary Figures S3–S21 show the direction of effects for each threat and conservation measure according to ordinal regression models, while model estimates are available in Supplementary Tables S1–S20.

**Figure 6.**
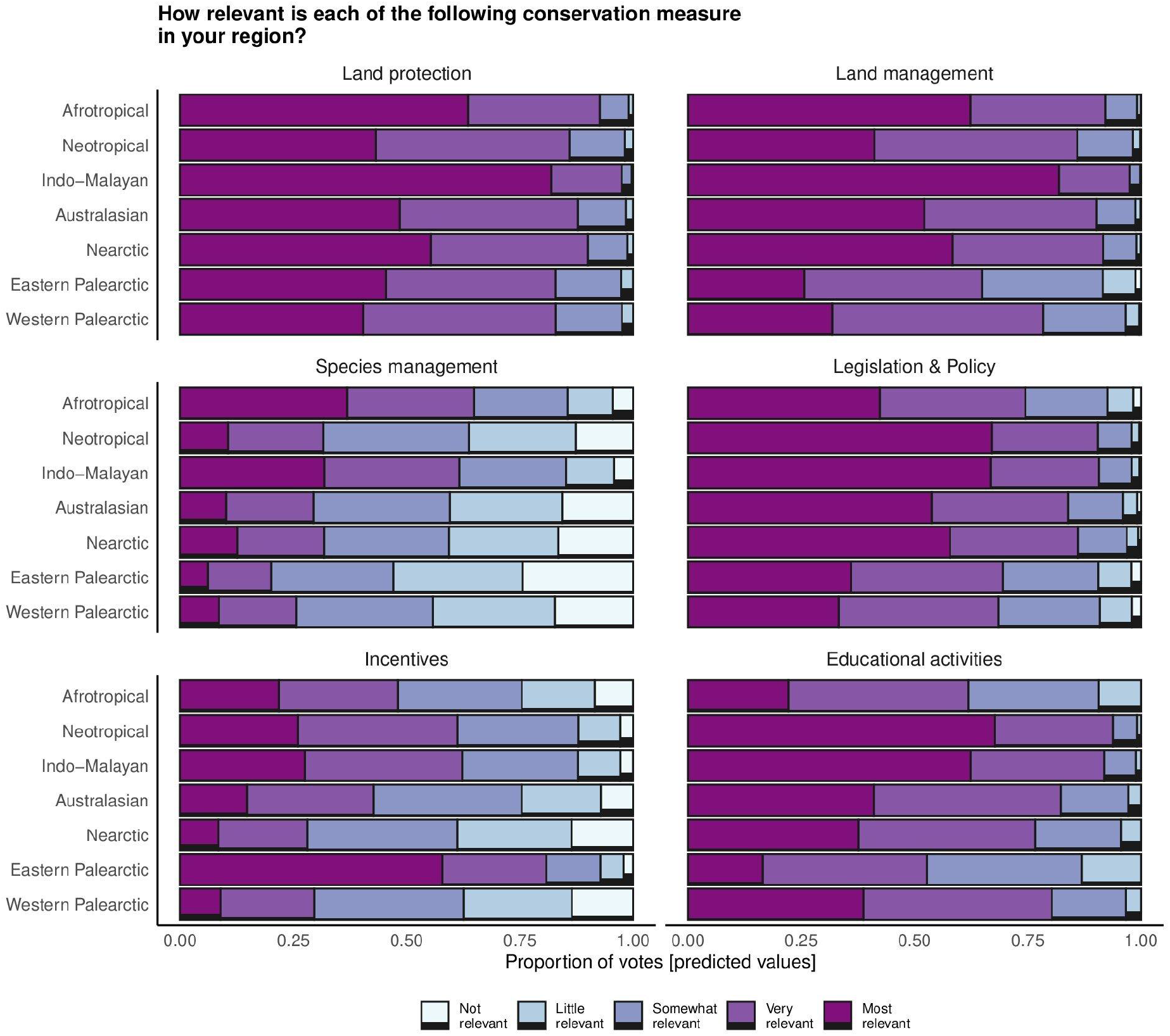
Importance of each conservation measure across biogeographic regions based on the responses by surveyed experts. Proportion of votes is expressed according to a Linkert scale. Note that scores are not the original values, but predictions from regression models that control for confounding factors related to respondents’ education and degree of experience with subterranean biology. Supplementary Figures S3–S21 show the direction of effects for each threat and conservation measure according to ordinal regression models, while model estimates are available in Supplementary Tables S1–S20.

## CONCLUSIONS

Subterranean ecosystems are impacted by multiple anthropogenic stressors whose effects remain to be fully quantified. Predictably, the degradation of subterranean ecosystems will result in irreplaceable loss of biodiversity and deprive societies of nature-based services, such as self-purifying groundwater processes, with high costs to human health and economies. Among the most pressing threats, experts identified habitat change and climate change. Legislation, land protection, and land management were perceived as the most effective actions to preserve the integrity of subterranean ecosystems, whereas species-level conservation was considered less important. Importantly, the relevance of threats and conservation varied across taxonomic groups and biogeographic regions, suggesting that conservation actions should be tailored on a case-by-case basis. Whenever hard data is lacking, expert opinion is a largely available, yet often underexploited source of information to identify threats and implement timely conservation of subterranean ecosystems and landscapes.

## Acknowledgements

A special thanks to all respondents in the online survey.

## Author contributions

VN and SM conceived the idea. All authors discussed the questionnaire structure. VN prepared the online survey, curated its dissemination, and cleaned the data. SM analyzed the data. VN and SM wrote the first draft. All authors contributed to the writing and analyses with suggestions and critical comments.

## Ethical statement

No data collection or scientific inquiries requiring ethics considerations were undertaken.

## Data availability statement

The database supporting the study is available in Figshare (upon acceptance). The R code to reproduce the analyses is available on GitHub (https://github.com/StefanoMammola/Analysis_Survey_subterranean_conservation) (link will start working upon acceptance).

## Authors’ Contributions

Conceptualization: SM, VN

Data collection: VN, SM

Data management: SM, VN

Data analysis: SM, EP, VN

Data visualization: SM, EP

Writing (first draft): VN, SM

Writing contributions: EP, PC, MI

All authors read the text, provided comments, suggestions and corrections, and approved the final version.

## Supplementary materials

Supplementary Text S1. Structure of the online survey in Google Form.

Figure S1–S21.

Table S1–S20

