## SUPPLEMENTARY MATERIAL for "An expert-based global assessment of threats and conservation measures for subterranean ecosystems"

Nanni V., et al.

### Table of content

Supplementary Text S1. Structure of the online survey in Google Form.

Table S1–S20

Figure S1–S21

**Supplementary Text S1.** Structure of the online survey in Google Form.

SECTION 1.

**1. What is your highest level of completed education?** [Single choice]

a. Primary/Secondary

b. University/College (Bachelor)

c. Master

d. PhD

**2. Indicate the option that best reflects your connection to subterranean ecosystems.** [Single choice]

a. Researcher/Scientist

b. Manager/Decision Maker

c. Recreational caver/ Speleologist

d. Educator

d. Other [Open-ended question]

**3. For how long have you held expertise on subterranean biology?** [Single choice]

a. Less than 5 years.

b. Between 5 and 10 years.

c. Between 10 and 20 years.

d. Over 20 years.

e. I don't have any expertise on subterranean ecosystems

**4. What biogeographic region does your expertise cover?** [Single choice]

a. Afrotropical

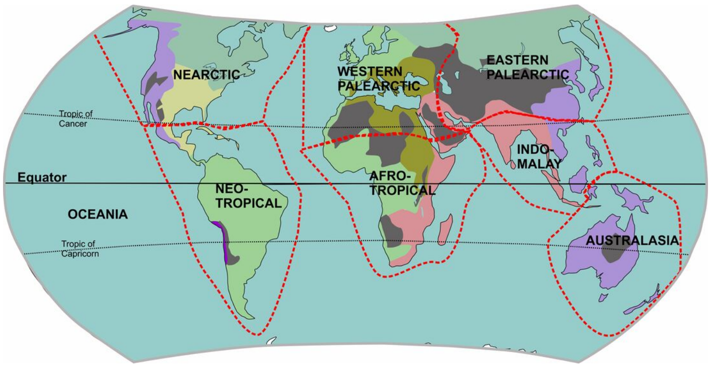


b. Australasian

c. Eastern Palearctic

d. Indo-Malayane.

e. Nearctic

f. Neotropical

g. Western Palearctic

**5. What type(s) of subterranean habitats does your expertise cover?** [Multiple choice]

a. Terrestrial (caves, lava tubes, MSS, mine, bunker, etc.)

b. Saltwater (marine cave, anchialine systems, etc.)

c. Freshwater (aquifer, groundwater, spring, flooded mine, water pipe, etc.)

SECTION 2.

**6. Does your expertise cover a specific group of subterranean organisms* (if your expertise cover more groups you will have the possibility to answer for these later in the questionnaire)?**
*** We refer here to native organisms. If you focus on alien/invasive species, these are considered a threat and so use 'none specific'.** [Single choice]

a. None specific [bring to SECTION 8]

b. Terrestrial invertebrates

c. Aquatic invertebrates

d. Cavefish

e. Bats

f. Terrestrial amphibians

g. Aquatic amphibians

h. Other vertebrates

i. Fungi

j. Prokaryotes/virus

k. Other photosynthetic organisms

**7. If you want to provide details about this specific group of subterranean organisms you focus on (e.g., "groundwater-dwelling Dytiscidae beetles", "all bats", "only the cave in my backyard", etc), please do so here.** [Open-ended question]

**8. How relevant is each of the following threats to this specific group of subterranean organisms (you may refer to direct and/or indirect threats)?** [Lickert scale]

| | not relevant | little relevance | somewhat relevant | very relevant | most relevant | unknown |
|---|---|---|---|---|---|---|
| Urbanisation | 1 | 2 | 3 | 4 | 5 | 6 |
| Agriculture/livestock farming/forestry | 1 | 2 | 3 | 4 | 5 | 6 |
| Recreational activities (e.g. tourism, recreational speleology) | 1 | 2 | 3 | 4 | 5 | 6 |
| Energy production & mining | 1 | 2 | 3 | 4 | 5 | 6 |
| Dams and water management | 1 | 2 | 3 | 4 | 5 | 6 |
| Wildfires | 1 | 2 | 3 | 4 | 5 | 6 |
| Pollution (e.g. pesticides, pharmaceuticals, plastic, and microplastic) | 1 | 2 | 3 | 4 | 5 | 6 |
| Invasive species | 1 | 2 | 3 | 4 | 5 | 6 |

| | | | | | | |
|---|---|---|---|---|---|---|
| Diseases (e.g. white nose syndrome, chithridiomycosis) | 1 | 2 | 3 | 4 | 5 | 6 |
| Climate change | 1 | 2 | 3 | 4 | 5 | 6 |
| Hunting/trapping/poaching | 1 | 2 | 3 | 4 | 5 | 6 |
| Natural geological events (e.g. volcanic eruptions, earthquakes) | 1 | 2 | 3 | 4 | 5 | 6 |
| Others | 1 | 2 | 3 | 4 | 5 | 6 |

**9. If you considered "other threats" as somewhat relevant or more in the previous question, please state these threats here. In your opinion, are there any other relevant future threats to this group of subterranean organisms not present in the previous question?** [Open-ended question]

**10. Thinking about the threats you consider the most relevant, can you describe how they impact this group of subterranean organisms?** [Open-ended question]

**11. How relevant is each of the following as conservation measures to this group of subterranean organisms you focus on?** [Lickert scale]

| | not relevant | little relevance | somewhat relevant | very relevant | most relevant | unknown |
|---|---|---|---|---|---|---|
| Land protection | 1 | 2 | 3 | 4 | 5 | 6 |
| Land management | 1 | 2 | 3 | 4 | 5 | 6 |
| Species management (e.g. harvest, trade, ex-situ conservation) | 1 | 2 | 3 | 4 | 5 | 6 |
| Education & awareness | 1 | 2 | 3 | 4 | 5 | 6 |
| Law & policy | 1 | 2 | 3 | 4 | 5 | 6 |
| Livelihood, economic & other incentives | 1 | 2 | 3 | 4 | 5 | 6 |
| Other protection measures | 1 | 2 | 3 | 4 | 5 | 6 |

**12. If you considered "Other protection measures" as somewhat relevant or more in the previous question, please state these here. In your opinion, are there any other relevant protection measures to this group of subterranean organisms not present in the previous question?** [Open-ended question]

**13. Does your expertise cover another group of subterranean organisms?** [Single scale]

a. Yes

b. No [bring to SECTION 3]

SECTION 3

**14. Please provide some references to published sources that support your answers if at all. Both references that you have authored or otherwise are acceptable and, if nothing else, personal communications as well.** [Open-ended question]

**15. If you have any other comment or suggestion, please feel free to write whatever you want in the text box below:** [Open-ended question]

**Supplementary Table S1. Estimated values for the Bayesian ordinal regression for 'Climate change'.** CI = confidence interval; ESS = Effective Sample Size; s.e. = Standard Error.

| *Climate change* | *Variable* | *Estimate* ± *s.e.* | **CI (95%)** | **Bulk_ESS** | **Tail_ESS** |
|---|---|---|---|---|---|
| Fixed Effects | Intercept[1] | -3.77 ± 1.44 | [-6.52, -0.85] | 1331 | 1748 |
| | Intercept[2] | -2.14 ± 1.43 | [-4.88, 0.78] | 1321 | 1703 |
| | Intercept[3] | -0.52 ± 1.42 | [-3.23, 2.43] | 1321 | 1717 |
| | Intercept[4] | 1.11 ± 1.42 | [-1.56, 4.12] | 1339 | 1629 |
| | Biogeography [Australasian] | -0.11 ± 0.95 | [-2.11, 1.69] | 1378 | 1120 |
| | Biogeography [Eastern Palearctic] | -0.73 ± 1.99 | [-4.51, 3.34] | 2813 | 2314 |
| | Biogeography [IndoMalayan] | -0.78 ± 1.08 | [-3.01, 1.31] | 1656 | 1697 |
| | Biogeography [Nearctic] | 0.44 ± 0.98 | [-1.61, 2.30] | 1407 | 1014 |
| | Biogeography [Neotropical] | -0.27 ± 0.95 | [-2.29, 1.51] | 1412 | 1139 |
| | Biogeography [Western Palearctic] | -0.54 ± 0.94 | [-2.48, 1.24] | 1369 | 1143 |
| | Taxon [Aquatic invertebrates] | 0.37 ± 0.59 | [-0.79, 1.53] | 1266 | 1901 |
| | Taxon [Bats] | -0.24 ± 0.67 | [-1.59, 1.03] | 1435 | 2076 |
| | Taxon [Cavefish] | 0.01 ± 0.75 | [-1.48, 1.46] | 1796 | 2160 |
| | Taxon [Microorganisms] | 0.15 ± 0.71 | [-1.25, 1.52] | 1871 | 2249 |
| | Taxon [None specific] | -0.41 ± 0.66 | [-1.70, 0.89] | 1716 | 1946 |
| | Taxon [Terrestrial vertebrates] | -2.71 ± 1.01 | [-4.74, -0.77] | 2311 | 2417 |
| | Taxon [Terrestrial invertebrates] | -0.50 ± 0.59 | [-1.66, 0.64] | 1284 | 2072 |
| *Random effects* | Education | 1.20 ± 0.81 | [0.32, 3.35] | 1442 | 1829 |
| | Profession | 0.97 ± 0.82 | [0.04, 3.04] | 865 | 1590 |

**Supplementary Table S2. Estimated values for the Bayesian ordinal regression for 'Geological events'.** CI = confidence interval; ESS = Effective Sample Size; s.e. = Standard Error.

| ***Geological events*** | ***Variable*** | ***Estimate ± s.e.*** | **CI (95%)** | **Bulk_ESS** | **Tail_ESS** |
|---|---|---|---|---|---|
| Fixed Effects | Intercept[1] | 1.69 ± 1.41 | [-0.97, 4.55] | 1238 | 1596 |
| | Intercept[2] | 2.97 ± 1.41 | [0.30, 5.81] | 1241 | 1691 |
| | Intercept[3] | 4.24 ± 1.42 | [1.55, 7.14] | 1255 | 1614 |
| | Intercept[4] | 5.51 ± 1.43 | [2.82, 8.46] | 1280 | 1615 |
| | Biogeography [Australasian] | 0.21 ± 0.97 | [-1.70, 2.16] | 1337 | 2124 |
| | Biogeography [Eastern Palearctic] | 0.11 ± 1.49 | [-2.90, 2.95] | 1818 | 2367 |
| | Biogeography [IndoMalayan] | 0.29 ± 1.07 | [-1.84, 2.49] | 1609 | 2429 |
| | Biogeography [Nearctic] | -0.83 ± 1.00 | [-2.79, 1.16] | 1276 | 2058 |
| | Biogeography [Neotropical] | -0.79 ± 0.99 | [-2.76, 1.21] | 1224 | 1844 |
| | Biogeography [Western Palearctic] | 0.27 ± 0.98 | [-1.71, 2.22] | 1240 | 1889 |
| | Taxon [Aquatic invertebrates] | 1.78 ± 0.93 | [0.09, 3.75] | 1175 | 1606 |
| | Taxon [Bats] | 1.76 ± 0.97 | [0.02, 3.78] | 1194 | 1590 |
| | Taxon [Cavefish] | 1.71 ± 1.01 | [-0.16, 3.82] | 1325 | 1557 |
| | Taxon [Microorganisms] | 3.27 ± 1.00 | [1.47, 5.38] | 1213 | 1794 |
| | Taxon [None specific] | 1.98 ± 1.00 | [0.14, 4.07] | 1229 | 1609 |
| | Taxon [Terrestrial vertebrates] | -0.59 ± 1.68 | [-4.37, 2.38] | 1683 | 2086 |
| | Taxon [Terrestrial invertebrates] | 1.44 ± 0.95 | [-0.28, 3.46] | 1119 | 1556 |
| *Random effects* | Education | 0.62 ± 0.55 | [0.03, 2.10] | 1466 | 1648 |
| | Profession | 0.64 ± 0.67 | [0.02, 2.41] | 1645 | 2226 |

**Supplementary Table S3. Estimated values for the Bayesian ordinal regression for 'Urbanisation'.** CI = confidence interval ; ESS = Effective Sample Size; s.e. = Standard Error.

| *Urbanisation* | *Variable* | *Estimate ± s.e.* | *CI (95%)* | *Bulk ESS* | *Tail ESS* |
|---|---|---|---|---|---|
| Fixed Effects | Intercept[1] | -0.75 ± 1.43 | [-3.43, 2.26] | 1316 | 1629 |
| | Intercept[2] | 0.80 ± 1.42 | [-1.89, 3.83] | 1301 | 1568 |
| | Intercept[3] | 2.34 ± 1.43 | [-0.30, 5.36] | 1295 | 1539 |
| | Intercept[4] | 3.88 ± 1.44 | [1.23, 6.91] | 1302 | 1583 |
| | Biogeography [Australasian] | -0.51 ± 1.03 | [-2.41, 1.56] | 1291 | 1683 |
| | Biogeography [Eastern Palearctic] | 3.11 ± 1.52 | [0.22, 6.20] | 1782 | 2201 |
| | Biogeography [IndoMalayan] | 1.84 ± 1.12 | [-0.29, 4.06] | 1463 | 2159 |
| | Biogeography [Nearctic] | 2.06 ± 1.05 | [0.13, 4.21] | 1308 | 1644 |
| | Biogeography [Neotropical] | 1.54 ± 1.03 | [-0.35, 3.67] | 1293 | 1768 |
| | Biogeography [Western Palearctic] | 1.12 ± 1.01 | [-0.73, 3.20] | 1223 | 1726 |
| | Taxon [Aquatic invertebrates] | 1.16 ± 0.54 | [0.12, 2.20] | 1386 | 2295 |
| | Taxon [Bats] | 0.76 ± 0.62 | [-0.47, 1.93] | 1597 | 2482 |
| | Taxon [Cavefish] | -0.29 ± 0.66 | [-1.61, 0.99] | 1673 | 2374 |
| | Taxon [Microorganisms] | 1.56 ± ***0.***65 | [0.25, 2.82] | 1776 | 2669 |
| | Taxon [None specific] | 1.58 ± ***0.***67 | [0.27, 2.92] | 1911 | 2757 |
| | Taxon [Terrestrial vertebrates] | -0.79 ± ***0.***79 | [-2.34, 0.71] | 2181 | 2508 |
| | Taxon [Terrestrial invertebrates] | 0.45 ± ***0.***54 | [-0.58, 1.51] | 1398 | 2344 |
| *Random effects* | Education | 0.77 ± 0.67 | [0.04, 2.59] | 1202 | 1714 |
| | Profession | 1.43 ± 0.80 | [0.44, 3.54] | 1397 | 1817 |

**Supplementary Table S4. Estimated values for the Bayesian ordinal regression for 'Agriculture *&* Forestry'.** CI = confidence interval; ESS = Effective Sample Size; s.e. = Standard Error.

| ***Agriculture & Forestry*** | ***Variable*** | ***Estimate* ± *s.e.*** | **CI (95%)** | **Bulk ESS** | **Tail ESS** |
|---|---|---|---|---|---|
| Fixed Effects | Intercept[1] | -3.19 ± 1.50 | [-6.01, -0.08] | 1326 | 1943 |
| | Intercept[2] | -1.33 ± 1.49 | [-4.12, 1.81] | 1309 | 1918 |
| | Intercept[3] | 0.53 ± 1.49 | [-2.24, 3.62] | 1330 | 1866 |
| | Intercept[4] | 2.38 ± 1.50 | [-0.36, 5.54] | 1343 | 1856 |
| | Biogeography [Australasian] | -0.53 ± 0.94 | [-2.32, 1.42] | 1125 | 1691 |
| | Biogeography [Eastern Palearctic] | 0.84 ± 1.53 | [-2.08, 3.93] | 1777 | 2444 |
| | Biogeography [IndoMalayan] | 1.87 ± 1.12 | [-0.25, 4.09] | 1568 | 2146 |
| | Biogeography [Nearctic] | -1.14 ± 0.97 | [-2.97, 0.83] | 1166 | 1667 |
| | Biogeography [Neotropical] | 1.57 ± 0.95 | [-0.20, 3.56] | 1118 | 1660 |
| | Biogeography [Western Palearctic] | 0.34 ± 0.92 | [-1.40, 2.25] | 1066 | 1395 |
| | Taxon [Aquatic invertebrates] | 0.55 ± 0.49 | [-0.40, 1.52] | 1489 | 2161 |
| | Taxon [Bats] | -0.17 ± 0.56 | [-1.28, 0.92] | 1852 | 2430 |
| | Taxon [Cavefish] | -0.72 ± 0.67 | [-2.05, 0.56] | 1937 | 2375 |
| | Taxon [Microorganisms] | 1.81 ± 0.65 | [0.57, 3.13] | 2061 | 2688 |
| | Taxon [None specific] | 0.58 ± 0.62 | [-0.62, 1.79] | 2067 | 2794 |
| | Taxon [Terrestrial vertebrates] | -2.68 ± 0.80 | [-4.23, -1.13] | 2484 | 2585 |
| | Taxon [Terrestrial invertebrates] | -0.67 ± 0.50 | [-1.65, 0.30] | 1599 | 2367 |
| *Random effects* | Education | 0.83 ± 0.68 | [0.09, 2.60] | 1128 | 1295 |
| | Profession | 1.85 ± 0.95 | [0.52, 4.20] | 1468 | 1670 |

**Supplementary Table S5. Estimated values for the Bayesian ordinal regression for 'Wildfires'.** CI = confidence interval; ESS = Effective Sample Size; s.e. = Standard Error.

| *Wildfires* | *Variable* | *Estimate ± s.e.* | **CI (95%)** | **Bulk_ ESS** | **Tail_ESS** |
|---|---|---|---|---|---|
| Fixed Effects | Intercept[1] | 3.11 ± 1.45 | [0.42, 6.08] | 1157 | 1580 |
| | Intercept[2] | 4.82 ± 1.47 | [2.12, 7.83] | 1152 | 1448 |
| | Intercept[3] | 6.53 ± 1.49 | [3.74, 9.56] | 1155 | 1511 |
| | Intercept[4] | 8.24 ± 1.52 | [5.38, 11.33] | 1168 | 1499 |
| | Biogeography [Australasian] | 2.22 ± 1.08 | [0.22, 4.40] | 1143 | 1758 |
| | Biogeography [Eastern Palearctic] | 2.32 ± 1.47 | [-0.43, 5.23] | 1485 | 2107 |
| | Biogeography [IndoMalayan] | 2.67 ± 1.16 | [0.46, 5.01] | 1299 | 1987 |
| | Biogeography [Nearctic] | 0.76 ± 1.10 | [-1.34, 2.99] | 1203 | 1892 |
| | Biogeography [Neotropical] | 2.19 ± 1.09 | [0.12, 4.37] | 1158 | 1831 |
| | Biogeography [Western Palearctic] | 1.59 ± 1.07 | [-0.43, 3.72] | 1126 | 1770 |
| | Taxon [Aquatic invertebrates] | 1.74 ± 0.79 | [0.28, 3.38] | 1224 | 1520 |
| | Taxon [Bats] | 3.52 ± 0.87 | [1.82, 5.32] | 1436 | 1689 |
| | Taxon [Cavefish] | 1.44 ± 0.89 | [-0.19, 3.26] | 1359 | 1913 |
| | Taxon [Microorganisms] | 2.49 ± 0.89 | [0.78, 4.28] | 1456 | 1719 |
| | Taxon [None specific] | 2.58 ± 0.83 | [1.06, 4.27] | 1307 | 1712 |
| | Taxon [Terrestrial vertebrates] | 3.08 ± 1.41 | [0.30, 5.90] | 2158 | 2434 |
| | Taxon [Terrestrial invertebrates] | 2.45 ± 0.80 | [0.96, 4.12] | 1291 | 1603 |
| *Random effects* | Education | 0.37 ± *0.41* | [0.01, 1.54] | 1706 | 1928 |
| | Profession | 1.05 ± *0.84* | [0.05, 3.23] | 1266 | 1858 |

**Supplementary Table S6. Estimated values for the Bayesian ordinal regression for 'Water management & Dams'.** CI = confidence interval; ESS = Effective Sample Size; s.e. = Standard Error.

| ***Water management & Dams*** | ***Variable*** | ***Estimate ± s.e.*** | **CI (95%)** | **Bulk_ESS** | **Tail_ESS** |
|---|---|---|---|---|---|
| Fixed Effects | Intercept[1] | -1.88 ± 1.21 | [-4.15, 0.53] | 1202 | 1779 |
| | Intercept[2] | -0.63 ± 1.20 | [-2.87, 1.78] | 1187 | 1796 |
| | Intercept[3] | 0.62 ± 1.20 | [-1.60, 3.01] | 1182 | 1829 |
| | Intercept[4] | 1.88 ± 1.20 | [-0.37, 4.27] | 1187 | 1883 |
| | Biogeography [Australasian] | 1.87 ± 1.03 | [-0.04, 3.96] | 1200 | 1941 |
| | Biogeography [Eastern Palearctic] | 6.42 ± 2.15 | [2.44, 11.07] | 2266 | 2373 |
| | Biogeography [IndoMalayan] | 1.28 ± 1.08 | [-0.78, 3.49] | 1330 | 2162 |
| | Biogeography [Nearctic] | 1.78 ± 1.04 | [-0.19, 3.90] | 1225 | 1955 |
| | Biogeography [Neotropical] | 2.39 ± 1.04 | [0.44, 4.50] | 1256 | 2091 |
| | Biogeography [Western Palearctic] | 1.65 ± 1.00 | [-0.23, 3.70] | 1174 | 1670 |
| | Taxon [Aquatic invertebrates] | 0.06 ± 0.49 | [-0.91, 1.03] | 1489 | 1932 |
| | Taxon [Bats] | -1.53 ± 0.59 | [-2.69, -0.37] | 1824 | 2014 |
| | Taxon [Cavefish] | -0.66 ± 0.63 | [-1.87, 0.60] | 1803 | 2128 |
| | Taxon [Microorganisms] | -0.51 ± 0.60 | [-1.64, 0.66] | 1949 | 1954 |
| | Taxon [None specific] | -0.95 ± 0.57 | [-2.03, 0.18] | 1840 | 2417 |
| | Taxon [Terrestrial vertebrates] | -4.50 ± 1.05 | [-6.73, -2.58] | 2740 | 2671 |
| | Taxon [Terrestrial invertebrates] | -1.11 ± 0.52 | [-2.14, -0.09] | 1483 | 2112 |
| Random effects | Education | -0.51 ± 0.60 | [-1.64, 0.66] | 1949 | 1954 |
| | Profession | -0.95 ± 0.57 | [-2.03, 0.18] | 1840 | 2417 |

**Supplementary Table S7. Estimated values for the Bayesian ordinal regression for 'Tourism *&* Recreation'.** CI = confidence interval; ESS = Effective Sample Size; s.e. = Standard Error.

| Tourism & Recreation | Variable | Estimate ± s.e. | CI (95%) | Bulk_ESS | Tail_ESS |
|---|---|---|---|---|---|
| Fixed Effects | Intercept[1] | -1.60 ± 1.15 | [-377, 0.62] | 1137 | 1891 |
| | Intercept[2] | 0.45 ± 1.14 | [-1.72, 2.69] | 1116 | 1881 |
| | Intercept[3] | 2.51 ± 1.15 | [0.35, 4.80] | 1108 | 2000 |
| | Intercept[4] | 4.56 ± 1.17 | [2.40, 6.91] | 1118 | 1968 |
| | Biogeography [Australasian] | -0.71 ± 1.01 | [-2.66, 1.30] | 1113 | 1544 |
| | Biogeography [Eastern Palearctic] | 2.26 ± 1.35 | [-0.42, 4.96] | 1538 | 2145 |
| | Biogeography [IndoMalayan] | 1.90 ± 1.12 | [-0.27, 4.09] | 1420 | 2080 |
| | Biogeography [Nearctic] | -0.84 ± 1.02 | [-2.83, 1.16] | 1138 | 1556 |
| | Biogeography [Neotropical] | 1.73 ± 1.02 | [-0.22, 3.75] | 1092 | 1538 |
| | Biogeography [Western Palearctic] | 1.14 ± 1.00 | [-0.80, 3.11] | 1162 | 1444 |
| | Taxon [Aquatic invertebrates] | 0.57 ± 0.52 | [-0.51, 1.56] | 1084 | 1764 |
| | Taxon [Bats] | 2.70 ± 0.62 | [1.49, 3.89] | 1301 | 2323 |
| | Taxon [Cavefish] | 0.76 ± ***0.***67 | [-0.54, 2.10] | 1322 | 2086 |
| | Taxon [Microorganisms] | 2.38 ± 0.63 | [1.13, 3.65] | 1328 | 2033 |
| | Taxon [None specific] | 0.97 ± 0.62 | [-0.28, 2.16] | 1348 | 1948 |
| | Taxon [Terrestrial vertebrates] | 1.13 ± 0.82 | [-0.47, 2.72] | 1841 | 2519 |
| | Taxon [Terrestrial invertebrates] | 1.47 ± 0.56 | [0.37, 2.54] | 1180 | 1751 |
| Random effects | Education | 2.38 ± ***0.***63 | [1.13, 3.65] | 1328 | 2033 |
| | Profession | 0.97 ± ***0.***62 | [-0.28, 2.16] | 1348 | 1948 |

**Supplementary Table S8. Estimated values for the Bayesian ordinal regression for 'Pollution'.** CI = confidence interval; ESS = Effective Sample Size; s.e. = Standard Error.

| *Pollution* | *Variable* | *Estimate ± s.e.* | **CI (95%)** | **Bulk_ESS** | **Tail_ESS** |
|---|---|---|---|---|---|
| Fixed Effects | Intercept[1] | -5.49 ± 1.49 | [-8.45, -2.62] | 1130 | 1586 |
| | Intercept[2] | -3.90 ± 1.47 | [-6.80, -1.06] | 1121 | 1499 |
| | Intercept[3] | -2.32 ± 1.46 | [-5.26, 0.55] | 1120 | 1504 |
| | Intercept[4] | -0.73 ± 1.46 | [-3.69, 2.11] | 1130 | 1454 |
| | Biogeography [Australasian] | -2.01 ± 1.16 | [-4.41, 0.12] | 1058 | 1541 |
| | Biogeography [Eastern Palearctic] | -2.53 ± 1.58 | [-5.66, 0.48] | 1389 | 1921 |
| | Biogeography [IndoMalayan] | -0.51 ± 1.19 | [-2.99, 1.69] | 1120 | 1948 |
| | Biogeography [Nearctic] | -1.00 ± 1.19 | [-3.43, 1.15] | 1052 | 1546 |
| | Biogeography [Neotropical] | 0.05 ± 1.17 | [-2.40, 2.19] | 1052 | 1594 |
| | Biogeography [Western Palearctic] | -0.43 ± 1.15 | [-2.78, 1.70] | 1014 | 1624 |
| | Taxon [Aquatic invertebrates] | -0.42 ± 0.62 | [-1.67, 0.76] | 1243 | 1799 |
| | Taxon [Bats] | -1.70 ± 0.69 | [-3.12, -0.39] | 1339 | 2215 |
| | Taxon [Cavefish] | -0.88 ± 0.78 | [-2.43, 0.61] | 1539 | 2097 |
| | Taxon [Microorganisms] | 0.85 ± 0.79 | [-0.73, 2.39] | 1644 | 2544 |
| | Taxon [None specific] | -0.99 ± 0 .70 | [-2.39, 0.43] | 1449 | 1985 |
| | Taxon [Terrestrial vertebrates] | -4.88 ± 1.38 | [-7.57, -2.12] | 3261 | 2724 |
| | Taxon [Terrestrial invertebrates] | -1.70 ± 0.63 | [-2.97, -0.52] | 1263 | 1985 |
| *Random effects* | Education | 1.04 ± 0.77 | [0.19, 3.02] | 1311 | 1279 |
| | Profession | 0.65 ± 0.61 | [0.03, 2.19] | 1434 | 1859 |

**Supplementary Table S9. Estimated values for the Bayesian ordinal regression for ‘Mining’.** CI = confidence interval; ESS = Effective Sample Size; s.e. = Standard Error.

| ***Mining*** | ***Variable*** | ***Estimate ± s.e.*** | **CI (95%)** | **Bulk_ESS** | **Tail_ESS** |
|---|---|---|---|---|---|
| Fixed Effects | Intercept[1] | 0.21 ± 1.37 | [-2.45, 2.91] | 1366 | 1576 |
| | Intercept[2] | 1.48 ± 1.37 | [-1.19, 4.17] | 1359 | 1745 |
| | Intercept[3] | 2.75 ± 1.37 | [0.06, 5.43] | 1360 | 1732 |
| | Intercept[4] | 4.02 ± 1.39 | [1.30, 6.73] | 1365 | 1716 |
| | Biogeography [Australasian] | 4.64 ± 1.03 | [2.72, 6.78] | 1327 | 1768 |
| | Biogeography [Eastern Palearctic] | 3.88 ± 1.92 | [0.26, 7.87] | 2083 | 2444 |
| | Biogeography [IndoMalayan] | 2.30 ± 1.02 | [0.33, 4.45] | 1417 | 1736 |
| | Biogeography [Nearctic] | 1.45 ± 0.98 | [-0.47, 3.48] | 1260 | 1780 |
| | Biogeography [Neotropical] | 2.93 ± 0.98 | [1.02, 4.99] | 1266 | 1773 |
| | Biogeography [Western Palearctic] | 2.32 ± 0.95 | [0.48, 4.34] | 1236 | 1740 |
| | Taxon [Aquatic invertebrates] | 0.16 ± 0.65 | [-1.10, 1.36] | 1082 | 1687 |
| | Taxon [Bats] | 0.58 ± 0.72 | [-0.86, 1.92] | 1181 | 2002 |
| | Taxon [Cavefish] | 0.66 ± 0.77 | [-0.85, 2.18] | 1401 | 2135 |
| | Taxon [Microorganisms] | 0.68 ± 0.74 | [-0.76, 2.11] | 1295 | 1996 |
| | Taxon [None specific] | -0.28 ± 0.71 | [-1.71, 1.08] | 1307 | 1937 |
| | Taxon [Terrestrial vertebrates] | 0.39 ± 1.00 | [-1.62, 2.31] | 1785 | 2405 |
| | Taxon [Terrestrial invertebrates] | 0.44 ± 0.67 | [-0.85, 1.75] | 1146 | 1727 |
| Random effects | Education | 0.79 ± 0.73 | [0.03, 2.69] | 885 | 1010 |
| | Profession | 1.12 ± ***0.90*** | [0.05, 3.36] | 940 | 1746 |

**Supplementary Table S10. Estimated values for the Bayesian ordinal regression for 'Invasive species'.** CI = confidence interval; ESS = Effective Sample Size; s.e. = Standard Error.

| **Invasive species** | **Variable** | **Estimate ± s.e.** | **CI (95%)** | **Bulk_ESS** | **Tail_ESS** |
|---|---|---|---|---|---|
| Fixed Effects | Intercept[1] | -1.12 ± 1.57 | [-4.06, 2.12] | 1210 | 1354 |
| | Intercept[2] | 0.26 ± 1.57 | [-2.68, 3.43] | 1216 | 1511 |
| | Intercept[3] | 1.64 ± 1.57 | [-1.27, 4.84] | 1228 | 1489 |
| | Intercept[4] | 3.02 ± 1.58 | [0.10, 6.23] | 1241 | 1541 |
| | Biogeography [Australasian] | 5.08 ± 1.07 | [-1.47, 2.71] | 972 | 1769 |
| | Biogeography [Eastern Palearctic] | 0.42 ± 1.53 | [-2.52, 3.48] | 1217 | 2405 |
| | Biogeography [IndoMalayan] | 0.81 ± 1.13 | [-1.45, 3.06] | 1129 | 2082 |
| | Biogeography [Nearctic] | 0.04 ± 1.08 | [-2.00, 2.21] | 1008 | 1741 |
| | Biogeography [Neotropical] | 1.48 ± 1.07 | [-0.55, 3.60] | 1004 | 1816 |
| | Biogeography [Western Palearctic] | 0.66 ± 1.07 | [-1.35, 2.79] | 990 | 1773 |
| | Taxon [Aquatic invertebrates] | -0.78 ± 0.52 | [-1.80, 0.26] | 1428 | 1748 |
| | Taxon [Bats] | -1.24 ± 0.61 | [-2.44, -0.07] | 1783 | 2278 |
| | Taxon [Cavefish] | -0.36 ± 0.67 | [-1.66, 0.95] | 1761 | 2203 |
| | Taxon [Microorganisms] | 0.02 ± 0.62 | [-1.20, 1.26] | 1851 | 2270 |
| | Taxon [None specific] | -0.18 ± 0.61 | [-1.40, 1.04] | 1939 | 2592 |
| | Taxon [Terrestrial vertebrates] | -3.04 ± 0.96 | [-4.96, -1.20] | 2235 | 2391 |
| | Taxon [Terrestrial invertebrates] | -0.49 ± 0.52 | [-1.53, 0.52] | 1546 | 1962 |
| Random effects | Education | 0.45 ± 0.47 | [0.02, 1.77] | 1536 | 1770 |
| | Profession | 1.95 ± 1.29 | [0.10, 5.00] | 861 | 776 |

**Supplementary Table S11. Estimated values for the Bayesian ordinal regression for 'Diseases'.** CI = confidence interval; ESS = Effective Sample Size; s.e. = Standard Error.

| ***Diseases*** | ***Variable*** | ***Estimate ± s.e.*** | **CI (95%)** | **Bulk_ESS** | **Tail_ESS** |
|---|---|---|---|---|---|
| Fixed Effects | Intercept[1] | -1.79 ± 1.31 | [-4.22, 0.87] | 1223 | 1792 |
| | Intercept[2] | -0.31 ± 1.31 | [-2.72, 2.35] | 1217 | 1750 |
| | Intercept[3] | 1.17 ± 1.31 | [-1.31, 3.85] | 1226 | 1746 |
| | Intercept[4] | 2.66 ± 1.33 | [0.14, 5.35] | 1246 | 1777 |
| | Biogeography [Australasian] | 0.71 ± 1.05 | [-1.33, 2.85] | 1128 | 1851 |
| | Biogeography [Eastern Palearctic] | -1.61 ± 1.55 | [-4.77, 1.35] | 1836 | 2584 |
| | Biogeography [IndoMalayan] | 2.12 ± 1.13 | [-0.08, 4.41] | 1321 | 1891 |
| | Biogeography [Nearctic] | 0.54 ± 1.11 | [-1.65, 2.74] | 1139 | 1730 |
| | Biogeography [Neotropical] | 0.25 ± 1.06 | [-1.83, 2.39] | 1125 | 1651 |
| | Biogeography [Western Palearctic] | 0.60 ± 1.06 | [-1.45, 2.71] | 1085 | 1664 |
| | Taxon [Aquatic invertebrates] | -2.06 ± 0.53 | [-3.11, -1.00] | 1694 | 2380 |
| | Taxon [Bats] | 0.73 ± 0.60 | [-0.42, 1.92] | 1819 | 2553 |
| | Taxon [Cavefish] | -1.09 ± 0.67 | [-2.45, 0.24] | 2092 | 2644 |
| | Taxon [Microorganisms] | -1.45 ± 0.66 | [-2.74, -0.14] | 2262 | 2992 |
| | Taxon [None specific] | 0.06 ± 0.61 | [-1.15, 1.26] | 2436 | 2839 |
| | Taxon [Terrestrial vertebrates] | 0.11 ± 0.96 | [-1.80, 1.99] | 2900 | 3064 |
| | Taxon [Terrestrial invertebrates] | -1.19 ± 0.53 | [-2.22, -0.11] | 1636 | 2455 |
| *Random effects* | Education | 0.92 ± 0.70 | [0.12, 2.66] | 1136 | 1017 |
| | Profession | 0.79 ± 0.59 | [0.05, 2.23] | 1135 | 1118 |

**Supplementary Table S12. Estimated values for the Bayesian ordinal regression for 'Poaching & Hunting'.** CI = confidence interval; ESS = Effective Sample Size; s.e. = Standard Error.

| *Poaching & Hunting* | *Variable* | *Estimate ± s.e.* | **CI (95%)** | **Bulk_ESS** | **Tail_ESS** |
|---|---|---|---|---|---|
| Fixed Effects | Intercept[1] | -2.46 ± 1.19 | [-4.77, -0.16] | 998 | 1860 |
| | Intercept[2] | -0.96 ± 1.18 | [-3.21, 1.31] | 1020 | 1831 |
| | Intercept[3] | 0.54 ± 1.18 | [-1.68, 2.84] | 1058 | 1842 |
| | Intercept[4] | 2.04 ± 1.19 | [-0.15, 4.33] | 1108 | 1872 |
| | Biogeography [Australasian] | -1.32 ± 0.96 | [-3.12, 0.56] | 936 | 1597 |
| | Biogeography [Eastern Palearctic] | -0.43 ± 1.35 | [-3.04, 2.29] | 1379 | 2214 |
| | Biogeography [IndoMalayan] | 1.06 ± 1.04 | [-0.92, 3.12] | 1212 | 1483 |
| | Biogeography [Nearctic] | -1.57 ± 0.97 | [-3.39, 0.31] | 947 | 1331 |
| | Biogeography [Neotropical] | -0.67 ± 0.95 | [-2.50, 1.25] | 943 | 1370 |
| | Biogeography [Western Palearctic] | -0.66 ± 0.94 | [-2.42, 1.24] | 909 | 1382 |
| | Taxon [Aquatic invertebrates] | -2.09 ± 0.59 | [-3.26, -0.94] | 1076 | 1888 |
| | Taxon [Bats] | -0.38 ± 0.63 | [-1.63, 0.85] | 1266 | 1814 |
| | Taxon [Cavefish] | -0.68 ± 0.69 | [-2.05, 0.63] | 1302 | 2296 |
| | Taxon [Microorganisms] | -1.89 ± 0.73 | [-3.32, -0.49] | 1359 | 2263 |
| | Taxon [None specific] | -0.77 ± 0.65 | [-2.04, 0.50] | 1329 | 2083 |
| | Taxon [Terrestrial vertebrates] | 0.24 ± 0.84 | [-1.44, 1.89] | 1761 | 2071 |
| | Taxon [Terrestrial invertebrates] | -0.72 ± 0.59 | [-1.90, 0.40] | 1070 | 1636 |
| *Random effects* | Education | 0.78 ± 0.65 | [0.06, 2.37] | 981 | 1459 |
| | Profession | 0.44 ± 0.43 | [0.02, 1.55] | 2101 | 2238 |

**Supplementary Table S13. Estimated values for the Bayesian ordinal regression for 'Other threats'.** CI = confidence interval; ESS = Effective Sample Size; s.e. = Standard Error.

| ***Other threats*** | ***Variable*** | ***Estimate ± s.e.*** | **CI (95%)** | **Bulk_ESS** | **Tail_ESS** |
|---|---|---|---|---|---|
| Fixed Effects | Intercept[1] | -1.02 ±1.42 | [-3.78, 1.87] | 1112 | 1585 |
| | Intercept[2] | -0.29 ± 1.42 | [-3.05, 2.58] | 1108 | 1537 |
| | Intercept[3] | 0.45 ± 1.42 | [-2.31, 3.34] | 1109 | 1571 |
| | Intercept[4] | 1.19 ± 1.43 | [-1.58, 4.12] | 1116 | 1706 |
| | Biogeography [Australasian] | 0.10 ± 1.13 | [-2.03, 2.43] | 1168 | 1784 |
| | Biogeography [Eastern Palearctic] | -1.89 ± 2.45 | [-7.05, 2.62] | 2156 | 2548 |
| | Biogeography [IndoMalayan] | 0.38 ± 1.19 | [-1.92, 2.86] | 1264 | 1557 |
| | Biogeography [Nearctic] | 0.09 ± 1.13 | [-2.05, 2.36] | 1054 | 1426 |
| | Biogeography [Neotropical] | -0.65 ± 1.13 | [-2.80, 1.72] | 1111 | 1343 |
| | Biogeography [Western Palearctic] | -0.06 ± 1.12 | [-2.19, 2.27] | 1082 | 1342 |
| | Taxon [Aquatic invertebrates] | 0.54 ± 0.75 | [-0.96, 2.05] | 1299 | 1361 |
| | Taxon [Bats] | -0.35 ± 0.81 | [-1.97, 1.25] | 1478 | 2050 |
| | Taxon [Cavefish] | -1.00 ± 0.85 | [-2.69, 0.65] | 1594 | 1996 |
| | Taxon [Microorganisms] | 0.35 ± 0.93 | [-1.47, 2.16] | 1908 | 2664 |
| | Taxon [None specific] | -0.44 ± 0.82 | [-2.02, 1.17] | 1549 | 1825 |
| | Taxon [Terrestrial vertebrates] | 1.11 ± 2.33 | [-3.58, 5.75] | 3295 | 2472 |
| | Taxon [Terrestrial invertebrates] | 0.43 ± 0.72 | [-0.98, 1.87] | 1337 | 1945 |
| *Random effects* | Education | 0.72 ± 0.63 | [0.04, 2.32] | 1558 | 1691 |
| | Profession | 0.88 ± 0.77 | [0.04, 2.82] | 1285 | 2235 |

**Supplementary Table S14. Estimated values for the Bayesian ordinal regression for 'Land protection'.** CI = confidence interval; ESS = Effective Sample Size; s.e. = Standard Error.

| ***Land protection*** | ***Variable*** | ***Estimate* ± *s.e.*** | **CI (95%)** | **Bulk_ESS** | **Tail_ESS** |
|---|---|---|---|---|---|
| Fixed Effects | Intercept[1] | -4.02 ± 1.48 | [-7.00, -1.17] | 1357 | 1703 |
| | Intercept[2] | -1.73 ± 1.46 | [-4.68, 1.10] | 1350 | 1610 |
| | Intercept[3] | 0.57 ± 1.46 | [-2.40, 3.44] | 1372 | 1645 |
| | Intercept[4] | -0.66 ± 1.08 | [-2.91, 1.30] | 1223 | 1584 |
| | Biogeography [Australasian] | -1.05 ± 1.67 | [-4.43, 2.27] | 1675 | 1990 |
| | Biogeography [Eastern Palearctic] | 1.05 ± 1.35 | [-1.68, 3.65] | 1765 | 2146 |
| | Biogeography [IndoMalayan] | -0.58 ± 1.10 | [-2.91, 1.38] | 1305 | 1524 |
| | Biogeography [Nearctic] | -1.18 ± 1.08 | [-3.42, 0.74] | 1315 | 1624 |
| | Biogeography [Neotropical] | -1.00 ± 1.07 | [-3.27, 0.92] | 1274 | 1521 |
| | Biogeography [Western Palearctic] | 0.36 ± 0.56 | [-0.73, 1.44] | 1432 | 2333 |
| | Taxon [Aquatic invertebrates] | 1.71 ± 0.67 | [0.37, 3.02] | 1769 | 2453 |
| | Taxon [Bats] | 0.50 ± 0.73 | [-0.93, 1.89] | 1810 | 2400 |
| | Taxon [Cavefish] | 0.58 ± 0.66 | [-0.71, 1.83] | 1593 | 2264 |
| | Taxon [Microorganisms] | 1.16 ± 0.69 | [-0.16, 2.53] | 1470 | 2219 |
| | Taxon [None specific] | 0.70 ± 0.89 | [-0.98, 2.51] | 2245 | 2671 |
| | Taxon [Terrestrial vertebrates] | 1.14 ± 0.58 | [0.01, 2.25] | 1443 | 2111 |
| | Taxon [Terrestrial invertebrates] | -4.02 ± 1.48 | [-7.00, -1.17] | 1357 | 1703 |
| *Random effects* | Education | 1.28 ± 0.79 | [0.41, 3.42] | 1337 | 2252 |
| | Profession | 0.99 ± 0.78 | [0.04, 2.93] | 906 | 1611 |

**Supplementary Table S15. Estimated values for the Bayesian ordinal regression for 'Land management'.** CI = confidence interval; ESS = Effective Sample Size; s.e. = Standard Error.

| *Land management* | *Variable* | *Estimate ± s.e.* | **CI (95%)** | **Bulk_ESS** | **Tail_ESS** |
|---|---|---|---|---|---|
| Fixed Effects | Intercept[1] | -6.11 ± 1.81 | [-9.48, -2.24] | 1354 | 1915 |
| | Intercept[2] | -3.77 ± 1.78 | [-7.11, -0.02] | 1327 | 1938 |
| | Intercept[3] | -1.44 ± 1.77 | [-4.77, 2.32] | 1313 | 1826 |
| | Intercept[4] | 0.89 ± 1.77 | [-2.42, 4.69] | 1310 | 1880 |
| | Biogeography [Australasian] | -0.41 ± 1.06 | [-2.45, 1.64] | 1178 | 1678 |
| | Biogeography [Eastern Palearctic] | -2.23 ± 1.69 | [-5.46, 1.04] | 1411 | 2132 |
| | Biogeography [IndoMalayan] | 1.13 ± 1.32 | [-1.34, 3.85] | 1535 | 2149 |
| | Biogeography [Nearctic] | -0.32 ± 1.08 | [-2.46, 1.77] | 1223 | 1789 |
| | Biogeography [Neotropical] | -0.85 ± 1.05 | [-2.88, 1.19] | 1153 | 1670 |
| | Biogeography [Western Palearctic] | -1.10 ± 1.03 | [-3.10, 0.89] | 1164 | 1688 |
| | Taxon [Aquatic invertebrates] | 0.67 ± 0.58 | [-0.48, 1.79] | 1306 | 2008 |
| | Taxon [Bats] | 0.75 ± 0.65 | [-0.54, 2.05] | 1579 | 2389 |
| | Taxon [Cavefish] | 1.08 ± 0.76 | [-0.39, 2.59] | 1921 | 2540 |
| | Taxon [Microorganisms] | 0.74 ± 0.68 | [-0.60, 2.08] | 1596 | 2070 |
| | Taxon [None specific] | 1.08 ± 0.71 | [-0.28, 2.48] | 1842 | 2455 |
| | Taxon [Terrestrial vertebrates] | -0.89 ± 0.90 | [-2.62, 0.83] | 2205 | 2618 |
| | Taxon [Terrestrial invertebrates] | 0.83 ± 0.59 | [-0.35, 1.97] | 1405 | 1961 |
| *Random effects* | Education | 1.35 ± 0.97 | [0.27, 3.77] | 1261 | 1553 |
| | Profession | 1.74 ± 1.21 | [0.12, 4.63] | 1014 | 1186 |

**Supplementary Table S16. Estimated values for the Bayesian ordinal regression for 'Species management'.** CI = confidence interval; ESS = Effective Sample Size; s.e. = Standard Error.

| *Species management* | *Variable* | *Estimate ± s.e.* | **CI (95%)** | **Bulk_ESS** | **Tail_ESS** |
|---|---|---|---|---|---|
| Fixed Effects | Intercept[1] | -3.19 ± 1.37 | [-5.84, -0.38] | 1171 | 1641 |
| | Intercept[2] | -1.71 ± 1.36 | [-4.35, 1.04] | 1170 | 1727 |
| | Intercept[3] | -0.24 ± 1.36 | [-2.87, 2.49] | 1180 | 1683 |
| | Intercept[4] | 1.24 ± 1.37 | [-1.39, 4.03] | 1196 | 1622 |
| | Biogeography [Australasian] | -0.91 ± 1.03 | [-2.94, 1.06] | 1165 | 1626 |
| | Biogeography [Eastern Palearctic] | -2.11 ± 1.39 | [-4.83, 0.57] | 1442 | 2130 |
| | Biogeography [IndoMalayan] | -0.07 ± 1.08 | [-2.26, 1.96] | 1217 | 1709 |
| | Biogeography [Nearctic] | -1.46 ± 1.04 | [-3.48, 0.51] | 1117 | 1512 |
| | Biogeography [Neotropical] | -1.07 ± 1.04 | [-3.15, 0.86] | 1132 | 1823 |
| | Biogeography [Western Palearctic] | -1.36 ± 1.01 | [-3.36, 0.57] | 1099 | 1585 |
| | Taxon [Aquatic invertebrates] | -0.60 ± 0.52 | [-1.64, 0.43] | 1447 | 1903 |
| | Taxon [Bats] | 0.15 ± 0.59 | [-1.00, 1.27] | 1670 | 2213 |
| | Taxon [Cavefish] | 1.08 ± 0.67 | [-0.21, 2.38] | 1629 | 2751 |
| | Taxon [Microorganisms] | -1.10 ± 0.65 | [-2.39, 0.18] | 1640 | 2206 |
| | Taxon [None specific] | -0.67 ± 0.60 | [-1.85, 0.50] | 1512 | 2151 |
| | Taxon [Terrestrial vertebrates] | -0.65 ± 0.83 | [-2.31, 0.95] | 2187 | 2757 |
| | Taxon [Terrestrial invertebrates] | 0.14 ± 0.52 | [-0.89, 1.16] | 1411 | 2169 |
| *Random effects* | Education | 0.54 ± 0.51 | [0.02, 1.91] | 1270 | 1411 |
| | Profession | 1.57 ± 0.74 | [0.64, 3.52] | 1690 | 2361 |

**Supplementary Table S17. Estimated values for the Bayesian ordinal regression for 'Legislation & Policy'.** CI = confidence interval; ESS = Effective Sample Size; s.e. = Standard Error.

| *Legislation & Policy* | *Variable* | *Estimate ± s.e.* | **CI (95%)** | **Bulk_ESS** | **Tail_ESS** |
|---|---|---|---|---|---|
| Fixed Effects | Intercept[1] | -3.50 ± 1.37 | [-6.15, -0.69] | 1080 | 1280 |
| | Intercept[2] | -1.79 ± 1.34 | [-4.33, 1.00] | 1055 | 1196 |
| | Intercept[3] | -0.09 ± 1.33 | [-2.60, 2.70] | 1056 | 1183 |
| | Intercept[4] | 1.62 ± 1.34 | [-0.90, 4.39] | 1068 | 1168 |
| | Biogeography [Australasian] | 0.54 ± 1.01 | [-1.43, 2.48] | 1184 | 1481 |
| | Biogeography [Eastern Palearctic] | 1.04 ± 1.36 | [-1.63, 3.67] | 1402 | 1949 |
| | Biogeography [IndoMalayan] | 1.08 ± 1.07 | [-1.06, 3.14] | 1343 | 1772 |
| | Biogeography [Nearctic] | 1.41 ± 1.03 | [-0.63, 3.44] | 1121 | 1376 |
| | Biogeography [Neotropical] | 1.64 ± 1.05 | [-0.42, 3.68] | 1167 | 1536 |
| | Biogeography [Western Palearctic] | 0.09 ± 1.00 | [-1.86, 2.01] | 1109 | 1385 |
| | Taxon [Aquatic invertebrates] | 0.99 ± 0.51 | [-0.05, 2.01] | 1499 | 2155 |
| | Taxon [Bats] | 0.09 ± 0.57 | [-1.07, 1.18] | 1650 | 2232 |
| | Taxon [Cavefish] | 0.61 ± 0.73 | [-0.79, 2.06] | 1975 | 2691 |
| | Taxon [Microorganisms] | 0.32 ± 0.64 | [-0.96, 1.56] | 2021 | 2632 |
| | Taxon [None specific] | -0.56 ± 0.63 | [-1.81, 0.65] | 1862 | 2527 |
| | Taxon [Terrestrial vertebrates] | -1.01 ± 0.76 | [-2.54, 0.51] | 2176 | 2710 |
| | Taxon [Terrestrial invertebrates] | -0.04 ± 0.52 | [-1.08, 0.95] | 1449 | 2247 |
| *Random effects* | Education | 0.93 ± 0.78 | [0.19, 2.90] | 918 | 1696 |
| | Profession | 0.53 ± 0.56 | [0.02, 1.86] | 1639 | 1837 |

**Supplementary Table S18. Estimated values for the Bayesian ordinal regression for 'Incentives'.** CI = confidence interval; ESS = Effective Sample Size; s.e. = Standard Error.

| ***Incentives*** | ***Variable*** | ***Estimate* ± *s.e.*** | **CI (95%)** | **Bulk_ESS** | **Tail_ESS** |
|---|---|---|---|---|---|
| Fixed Effects | Intercept[1] | -2.27 ± 1.52 | [-5.22, 0.77] | 1116 | 1732 |
| | Intercept[2] | -0.69 ± 1.52 | [-3.64, 2.31] | 1106 | 1749 |
| | Intercept[3] | 0.89 ± 1.51 | [-2.04, 3.88] | 1107 | 1741 |
| | Intercept[4] | 2.46 ± 1.52 | [-0.47, 5.46] | 1114 | 1741 |
| | Biogeography [Australasian] | -0.07 ± 1.26 | [-2.52, 2.43] | 1063 | 1814 |
| | Biogeography [Eastern Palearctic] | 2.18 ± 1.70 | [-1.02, 5.57] | 1306 | 2150 |
| | Biogeography [IndoMalayan] | 0.56 ± 1.30 | [-1.92, 3.12] | 1110 | 1931 |
| | Biogeography [Nearctic] | -1.19 ± 1.28 | [-3.66, 1.35] | 976 | 1574 |
| | Biogeography [Neotropical] | 0.46 ± 1.26 | [-1.94, 2.93] | 954 | 1524 |
| | Biogeography [Western Palearctic] | -0.81 ± 1.26 | [-3.24, 1.64] | 954 | 1411 |
| | Taxon [Aquatic invertebrates] | 0.62 ± 0.52 | [-0.41, 1.65] | 1520 | 2405 |
| | Taxon [Bats] | 0.80 ± 0.58 | [-0.33, 1.96] | 1856 | 2634 |
| | Taxon [Cavefish] | 0.87 ± 0.65 | [-0.46, 2.11] | 1836 | 2556 |
| | Taxon [Microorganisms] | 1.08 ± 0.59 | [-0.04, 2.26] | 1742 | 2255 |
| | Taxon [None specific] | 0.67 ± 0.57 | [-0.44, 1.81] | 1865 | 2492 |
| | Taxon [Terrestrial vertebrates] | -2.08 ± 1.10 | [-4.45, -0.04] | 3360 | 2876 |
| | Taxon [Terrestrial invertebrates] | 0.53 ± 0.53 | [-0.50, 1.55] | 1465 | 2247 |
| *Random effects* | Education | 1.13 ± 0.72 | [0.30, 3.07] | 1573 | 2355 |
| | Profession | 0.84 ± 0.63 | [0.05, 2.38] | 1374 | 1595 |

**Supplementary Table S19. Estimated values for the Bayesian ordinal regression for 'Educational activities'.** CI = confidence interval; ESS = Effective Sample Size; s.e. = Standard Error.

| **Educational activities** | ***Variable*** | ***Estimate* ± *s.e.*** | **CI (95%)** | **Bulk_ESS** | **Tail_ESS** |
|---|---|---|---|---|---|
| Fixed Effects | Intercept[1] | -2.14 ± 1.53 | [-5.06, 1.07] | 1237 | 1677 |
| | Intercept[2] | -0.06 ± 1.53 | [-2.95, 3.19] | 1207 | 1688 |
| | Intercept[3] | 2.02 ± 1.54 | [-0.90, 5.30] | 1201 | 1790 |
| | Intercept[4] | 0.79 ± 1.02 | [-1.26, 2.72] | 1269 | 1960 |
| | Biogeography [Australasian] | 0.02 ± 1.41 | [-2.77, 2.70] | 1680 | 2338 |
| | Biogeography [Eastern Palearctic] | 1.95 ± 1.10 | [-0.28, 4.12] | 1384 | 1972 |
| | Biogeography [IndoMalayan] | -0.04 ± 1.04 | [-2.18, 1.96] | 1274 | 1911 |
| | Biogeography [Nearctic] | 2.13 ± 1.04 | [0.09, 4.14] | 1194 | 1933 |
| | Biogeography [Neotropical] | 0.71 ± 1.01 | [-1.31, 2.59] | 1111 | 1784 |
| | Biogeography [Western Palearctic] | 0.62 ± 0.55 | [-0.46, 1.69] | 1298 | 1862 |
| | Taxon [Aquatic invertebrates] | 0.92 ± 0.62 | [-0.36, 2.14] | 1505 | 1950 |
| | Taxon [Bats] | 0.29 ± 0.69 | [-1.08, 1.61] | 1700 | 2173 |
| | Taxon [Cavefish] | 1.84 ± 0.65 | [0.55, 3.10] | 1546 | 2479 |
| | Taxon [Microorganisms] | 1.21 ± 0.66 | [-0.11, 2.49] | 1479 | 1916 |
| | Taxon [None specific] | 0.55 ± 0.83 | [-1.09, 2.17] | 2247 | 2459 |
| | Taxon [Terrestrial vertebrates] | 0.49 ± 0.55 | [-0.62, 1.57] | 1363 | 1968 |
| | Taxon [Terrestrial invertebrates] | -2.14 ± 1.53 | [-5.06, 1.07] | 1237 | 1677 |
| *Random effects* | Education | 0.96 ± 0.79 | [0.05, 3.02] | 1102 | 1475 |
| | Profession | 1.33 ± 0.83 | [0.25, 3.43] | 1336 | 1264 |

**Supplementary Table S20. Estimated values for the Bayesian ordinal regression for 'Other protection'.** CI = confidence interval; ESS = Effective Sample Size; s.e. = Standard Error.

| ***Other protection*** | ***Variable*** | ***Estimate ± s.e.*** | **CI (95%)** | **Bulk_ESS** | **Tail_ESS** |
|---|---|---|---|---|---|
| Fixed Effects | Intercept[1] | -2.80 ± 1.56 | [-6.00, 0.16] | 1274 | 1373 |
| | Intercept[2] | -1.90 ± 1.55 | [-5.07, 1.04] | 1271 | 1406 |
| | Intercept[3] | -0.99 ± 1.55 | [-4.16, 1.93] | 1272 | 1346 |
| | Intercept[4] | -0.08 ± 1.55 | [-3.26, 2.82] | 1276 | 1555 |
| | Biogeography [Australasian] | -1.36 ± 1.10 | [-3.55, 0.77] | 1179 | 1760 |
| | Biogeography [Eastern Palearctic] | -1.10 ± 1.59 | [-4.29, 1.93] | 1631 | 2511 |
| | Biogeography [IndoMalayan] | -2.01 ± 1.20 | [-4.37, 0.35] | 1295 | 1723 |
| | Biogeography [Nearctic] | -1.26 ± 1.14 | [-3.47, 0.89] | 1078 | 1793 |
| | Biogeography [Neotropical] | -0.84 ± 1.09 | [-3.04, 1.18] | 1180 | 1854 |
| | Biogeography [Western Palearctic] | -0.66 ± 1.11 | [-2.92, 1.42] | 1080 | 1792 |
| | Taxon [Aquatic invertebrates] | 0.27 ± 0.72 | [-1.19, 1.69] | 1398 | 1821 |
| | Taxon [Bats] | -0.93 ± 0.82 | [-2.60, 0.64] | 1594 | 2169 |
| | Taxon [Cavefish] | -0.92 ± 0.86 | [-2.61, 0.73] | 1437 | 2051 |
| | Taxon [Microorganisms] | -0.03 ± 0.80 | [-1.64, 1.53] | 1710 | 2341 |
| | Taxon [None specific] | 0.59 ± 0.84 | [-1.10, 2.25] | 1669 | 2207 |
| | Taxon [Terrestrial vertebrates] | -46.774 ± 49.520 | [-16.1622, -22.40] | 1138 | 570 |
| | Taxon [Terrestrial invertebrates] | -0.78 ± 0.73 | [-2.26, 0.64] | 1493 | 1864 |
| *Random effects* | Education | 0.96 ± 0.75 | [0.08, 2.92] | 1175 | 1185 |
| | Profession | 1.47 ± 0.94 | [0.15, 3.81] | 1177 | 1228 |

**Supplementary Figures S1.** Comparison between observed scores (proportion of votes for the 5 score levels) and predicted scores (predicted probability for the 5 score levels) for the different threats considered in the survey. Predicted values are based on Bayesian ordinal regression models.

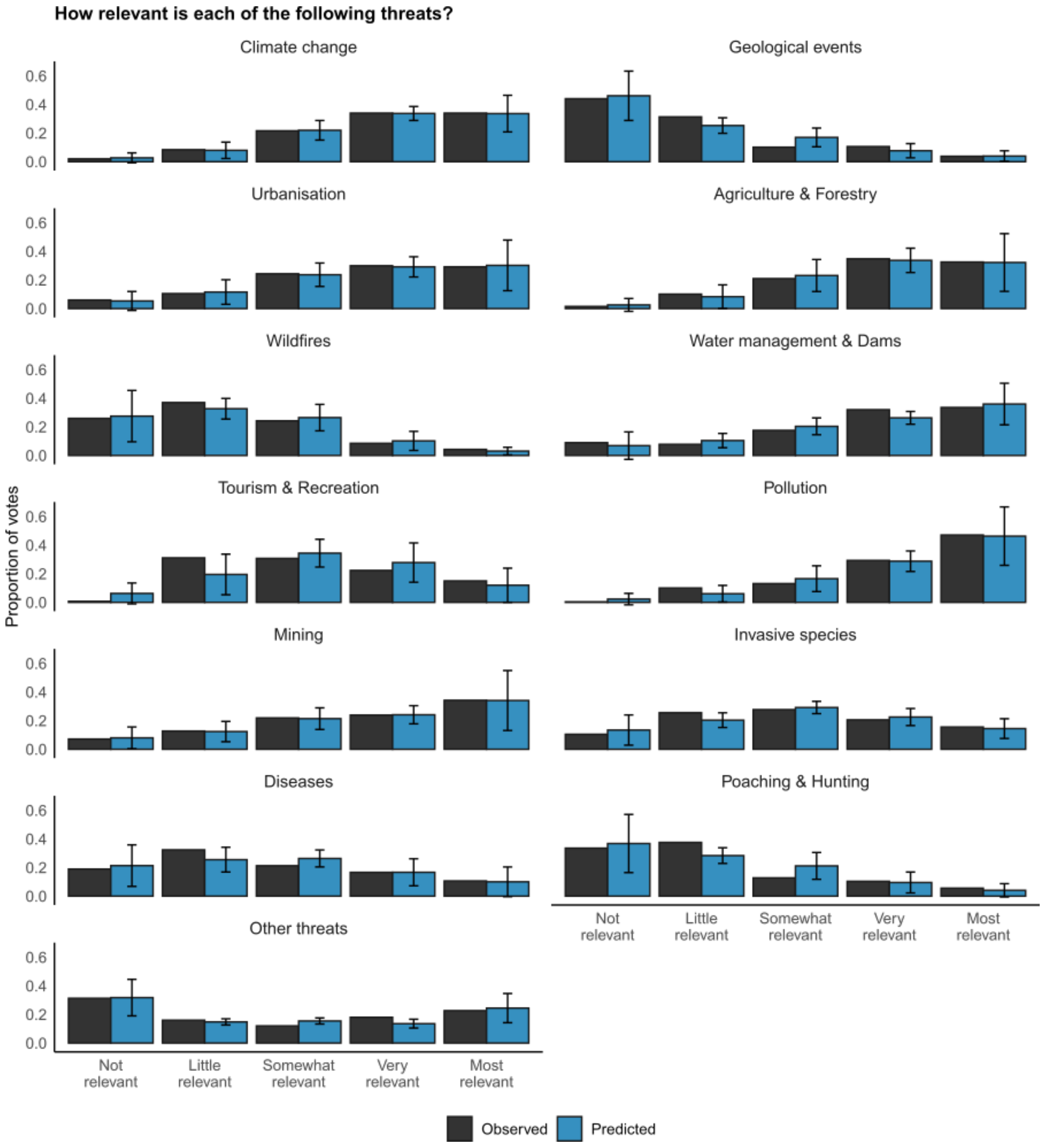

**Supplementary Figures S2.** Comparison between observed scores (proportion of votes for the 5 score levels) and predicted scores (predicted probability for the 5 score levels) for the different conservation measures considered in the survey. Predicted values are based on Bayesian ordinal regression models.

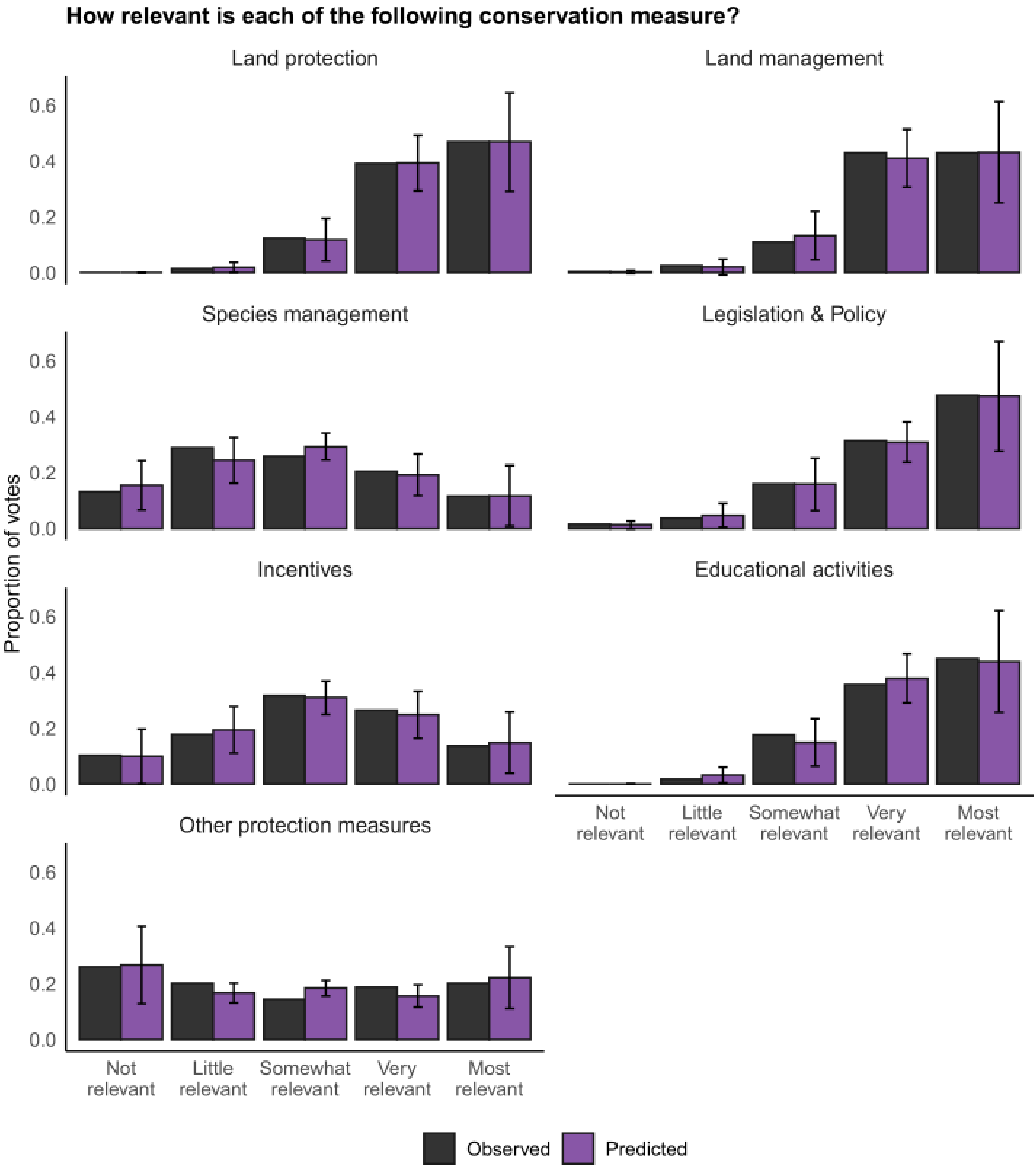

**Supplementary Figure S3**. Results of the Bayesian ordinal regression for ‘Climate change’. Bayesian model density plots computed from posterior draws, with uncertainty intervals shown as shaded areas under the curves. Probability mass corresponds to 90%.

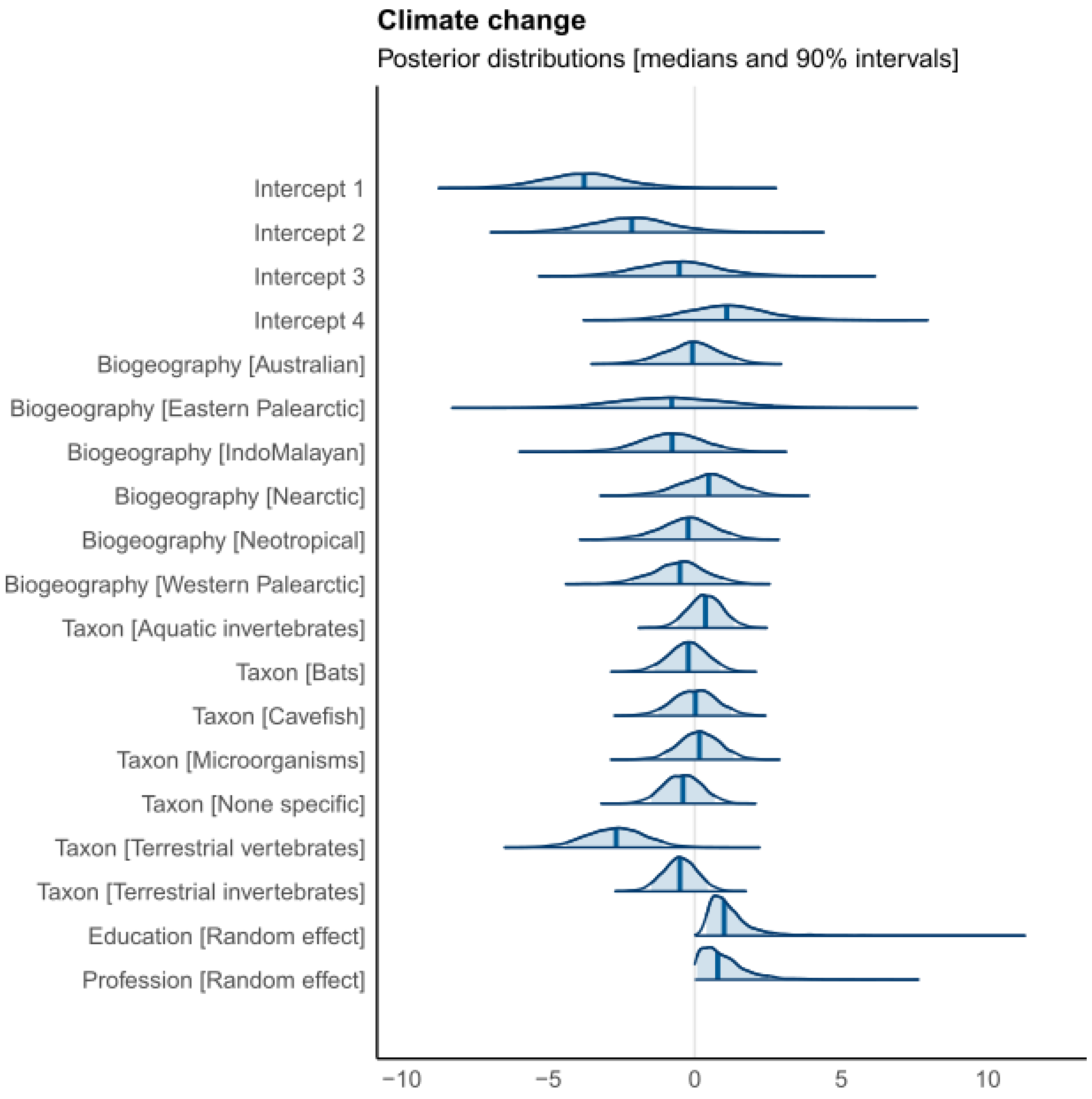

**Supplementary Figure S4**. Results of the Bayesian ordinal regression for ‘Geological events’. Bayesian model density plots computed from posterior draws, with uncertainty intervals shown as shaded areas under the curves. Probability mass corresponds to 90%.

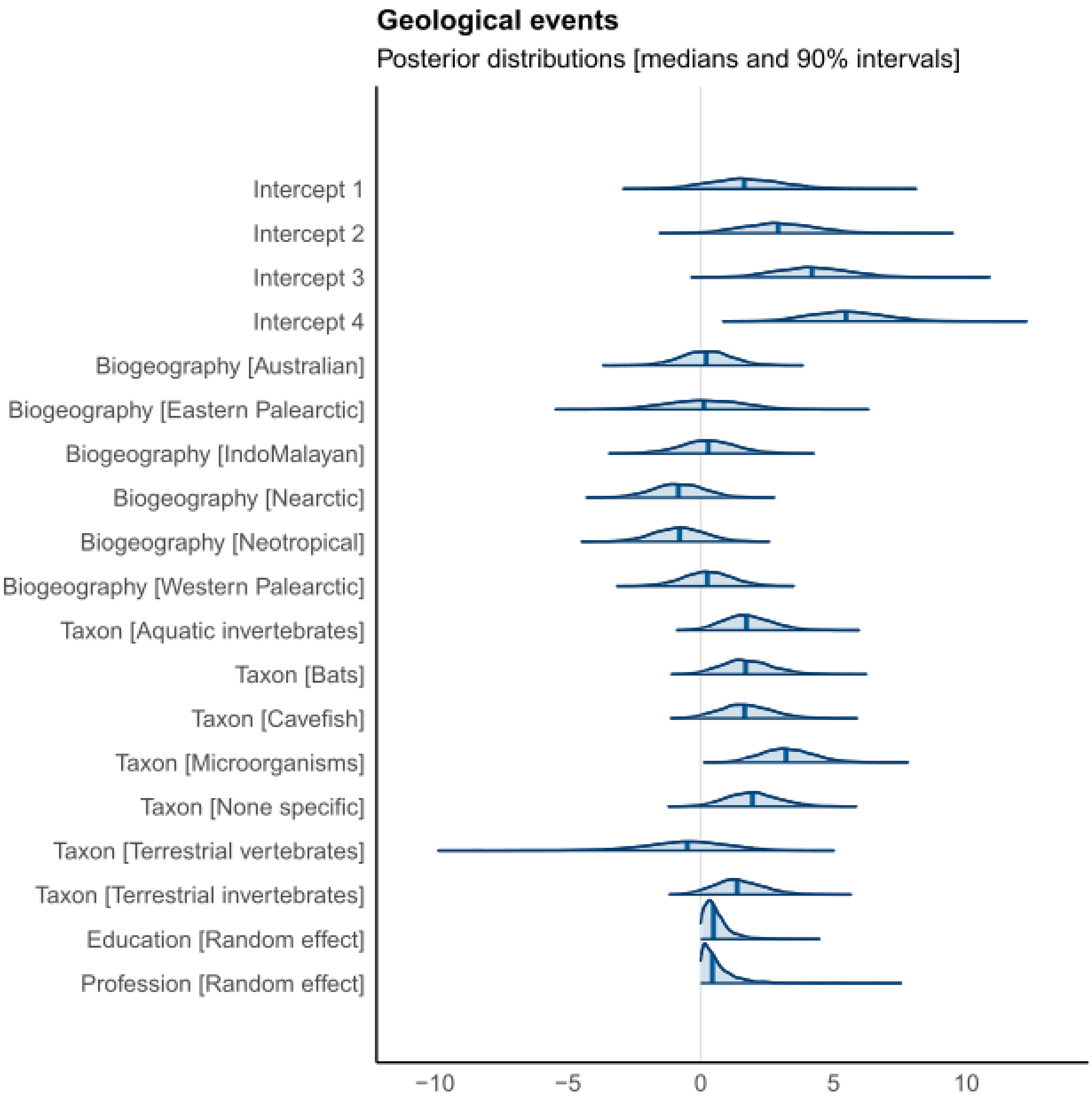

**Supplementary Figure S5**. Results of the Bayesian ordinal regression for ‘Urbanisation’. Bayesian model density plots computed from posterior draws, with uncertainty intervals shown as shaded areas under the curves. Probability mass corresponds to 90%.

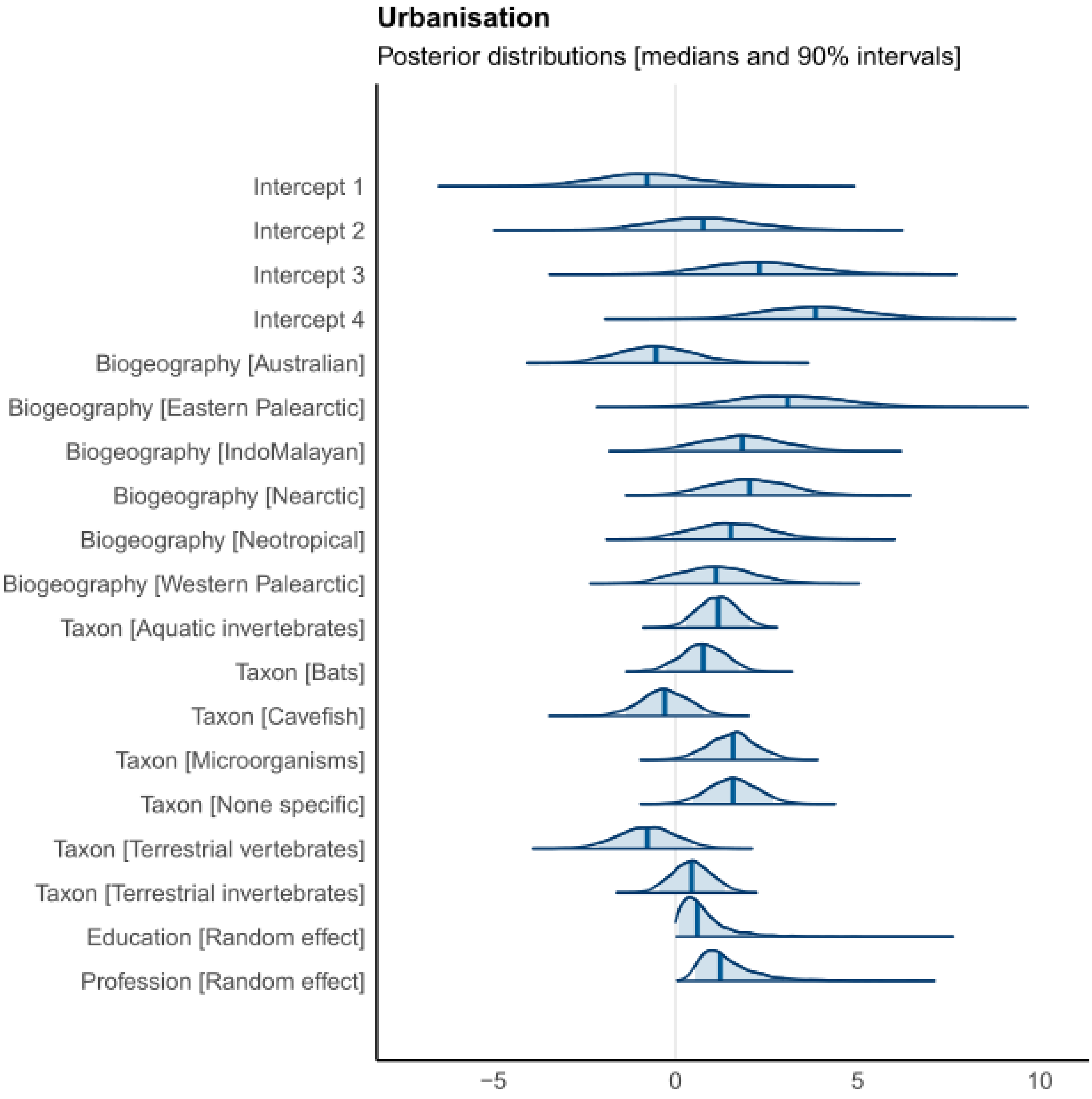

**Supplementary Figure S6**. Results of the Bayesian ordinal regression for 'Agriculture & Forestry'. Bayesian model density plots computed from posterior draws, with uncertainty intervals shown as shaded areas under the curves. Probability mass corresponds to 90%.

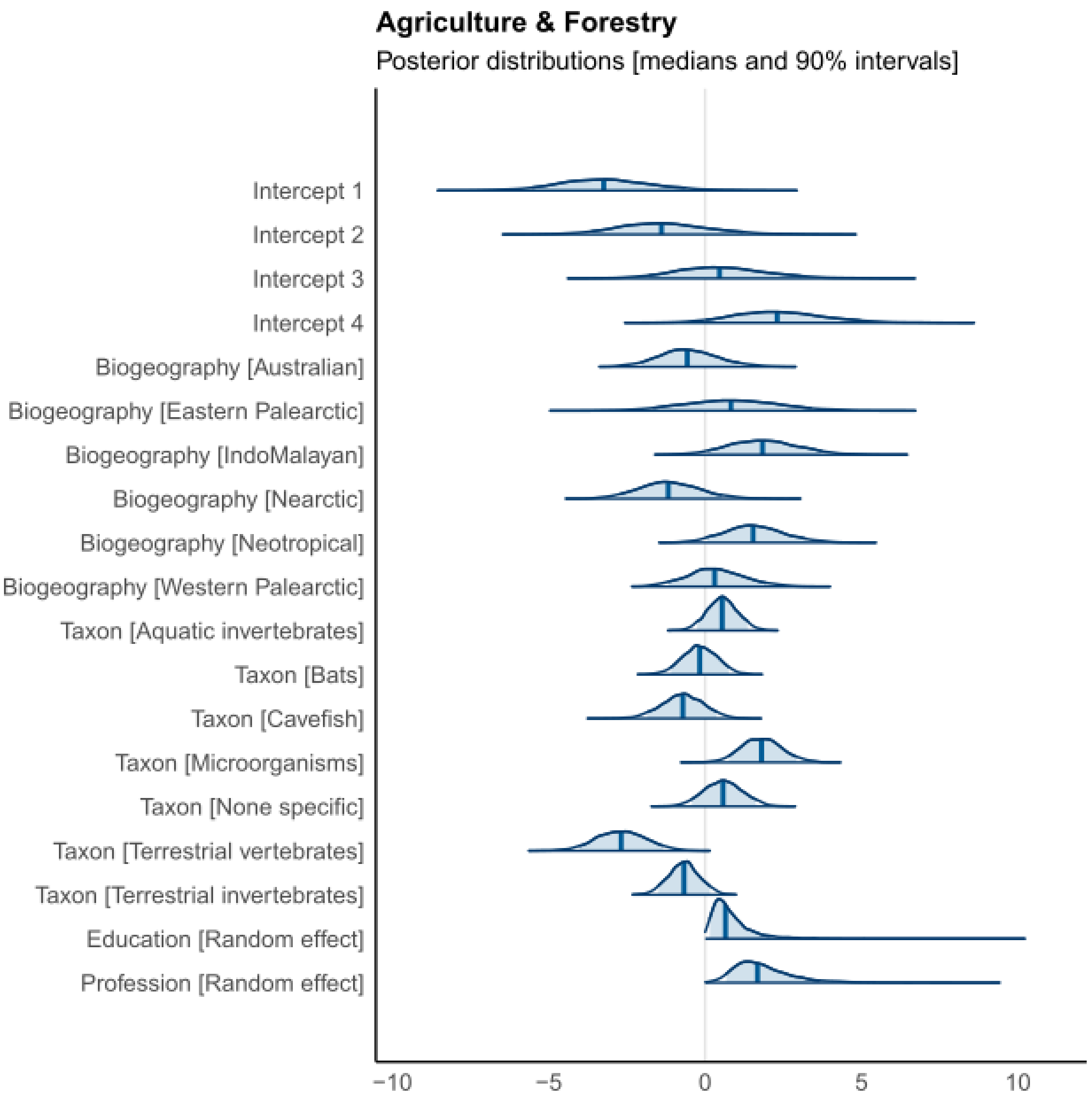

**Supplementary Figure S7**. Results of the Bayesian ordinal regression for 'Wildfires'. Bayesian model density plots computed from posterior draws, with uncertainty intervals shown as shaded areas under the curves. Probability mass corresponds to 90%.

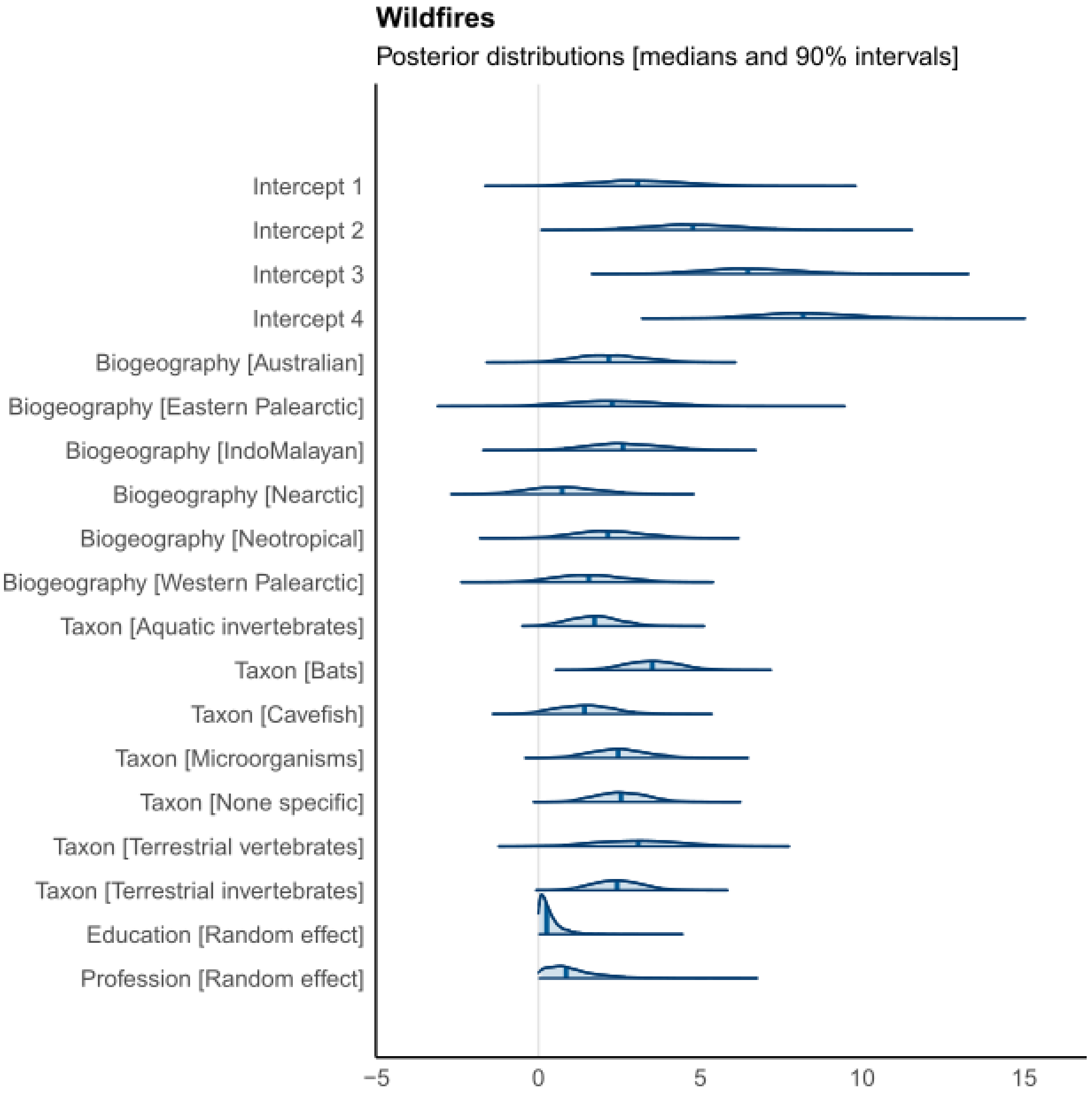

**Supplementary Figure S8**. Results of the Bayesian ordinal regression for ‘Water management & Dams’. Bayesian model density plots computed from posterior draws, with uncertainty intervals shown as shaded areas under the curves. Probability mass corresponds to 90%.

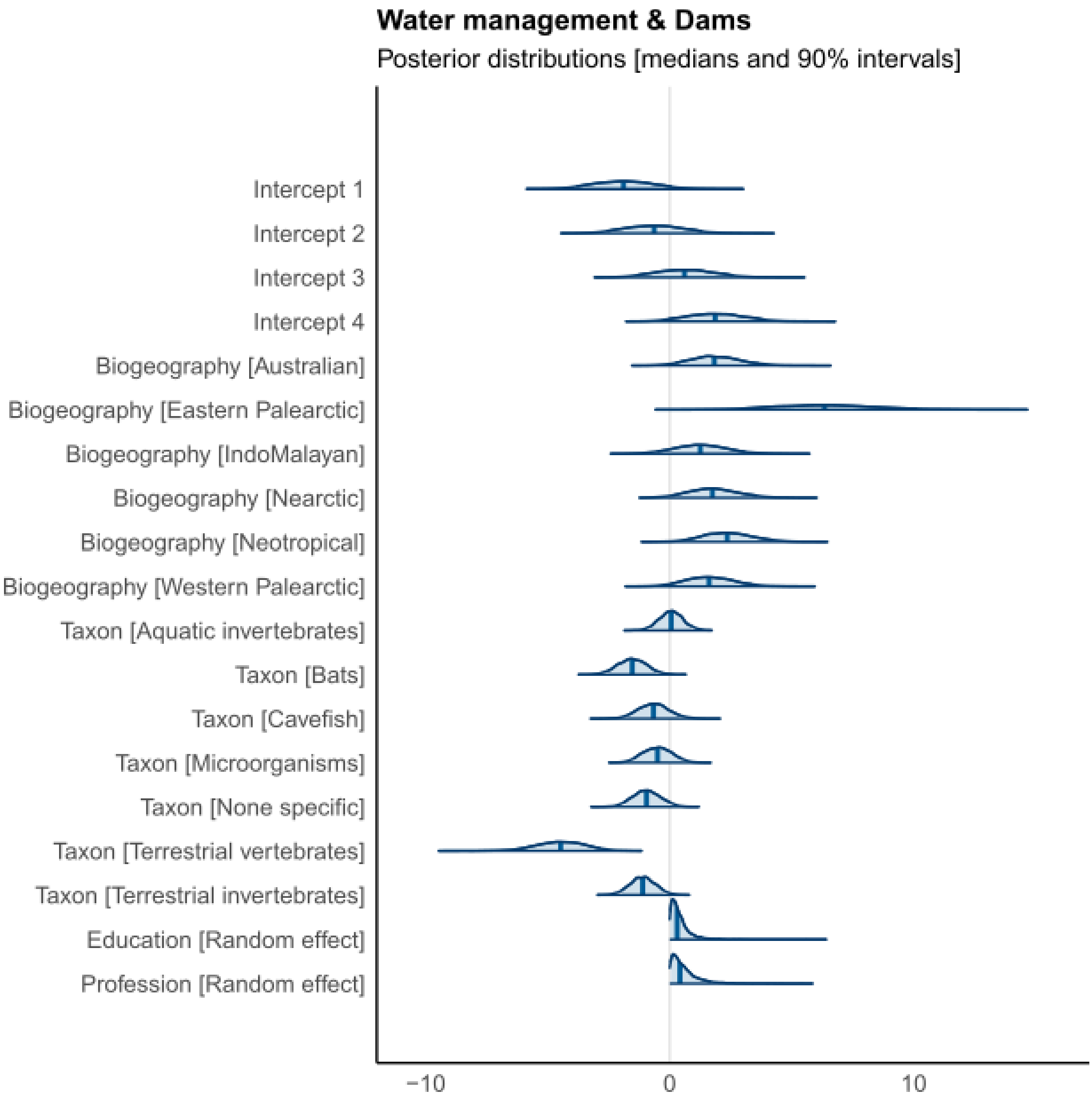

**Supplementary Figure S9**. Results of the Bayesian ordinal regression for ‘Tourism & Recreation’. Bayesian model density plots computed from posterior draws, with uncertainty intervals shown as shaded areas under the curves. Probability mass corresponds to 90%.

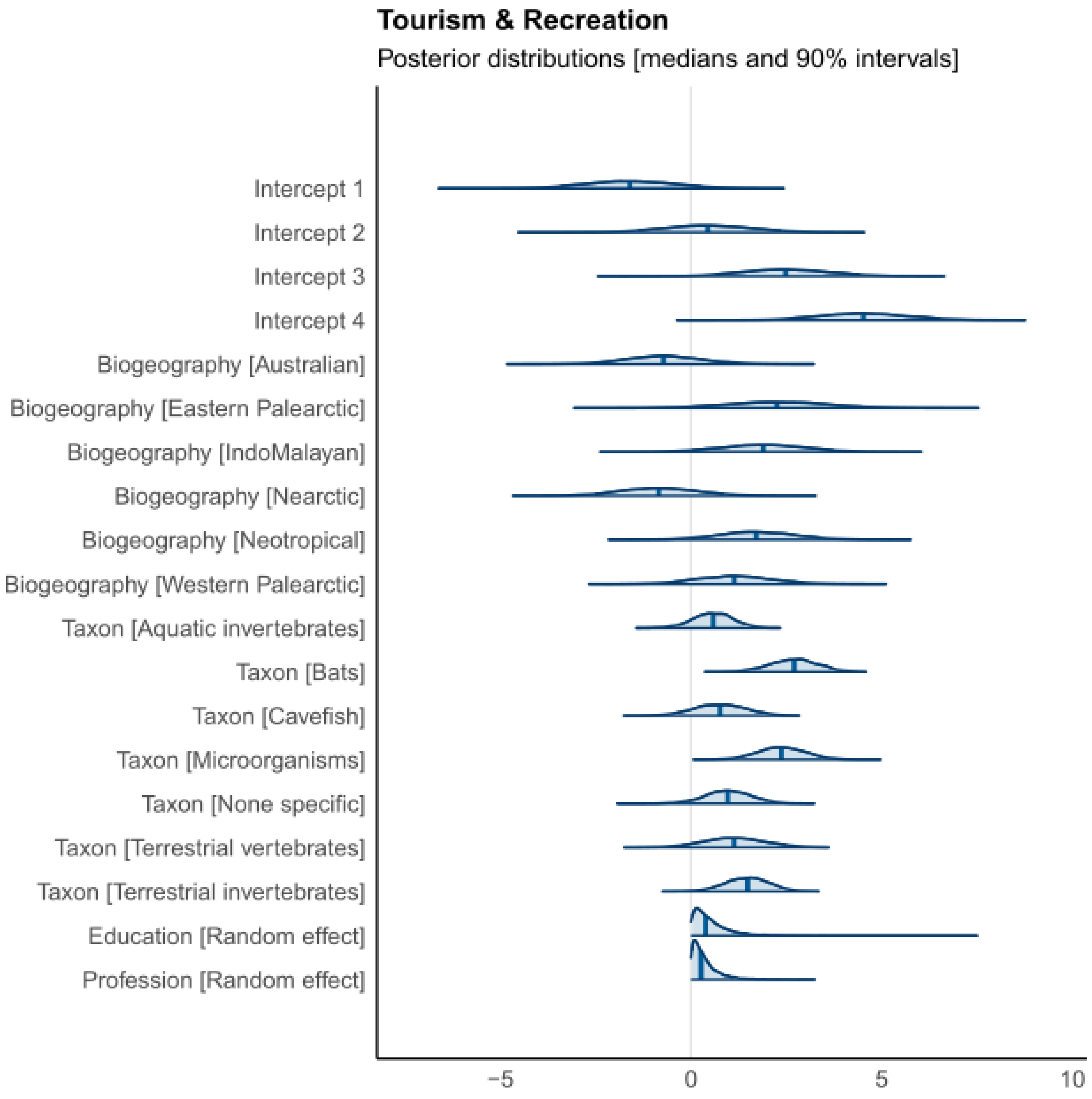

**Supplementary Figure S10**. Results of the Bayesian ordinal regression for ‘Pollution’. Bayesian model density plots computed from posterior draws, with uncertainty intervals shown as shaded areas under the curves. Probability mass corresponds to 90%.

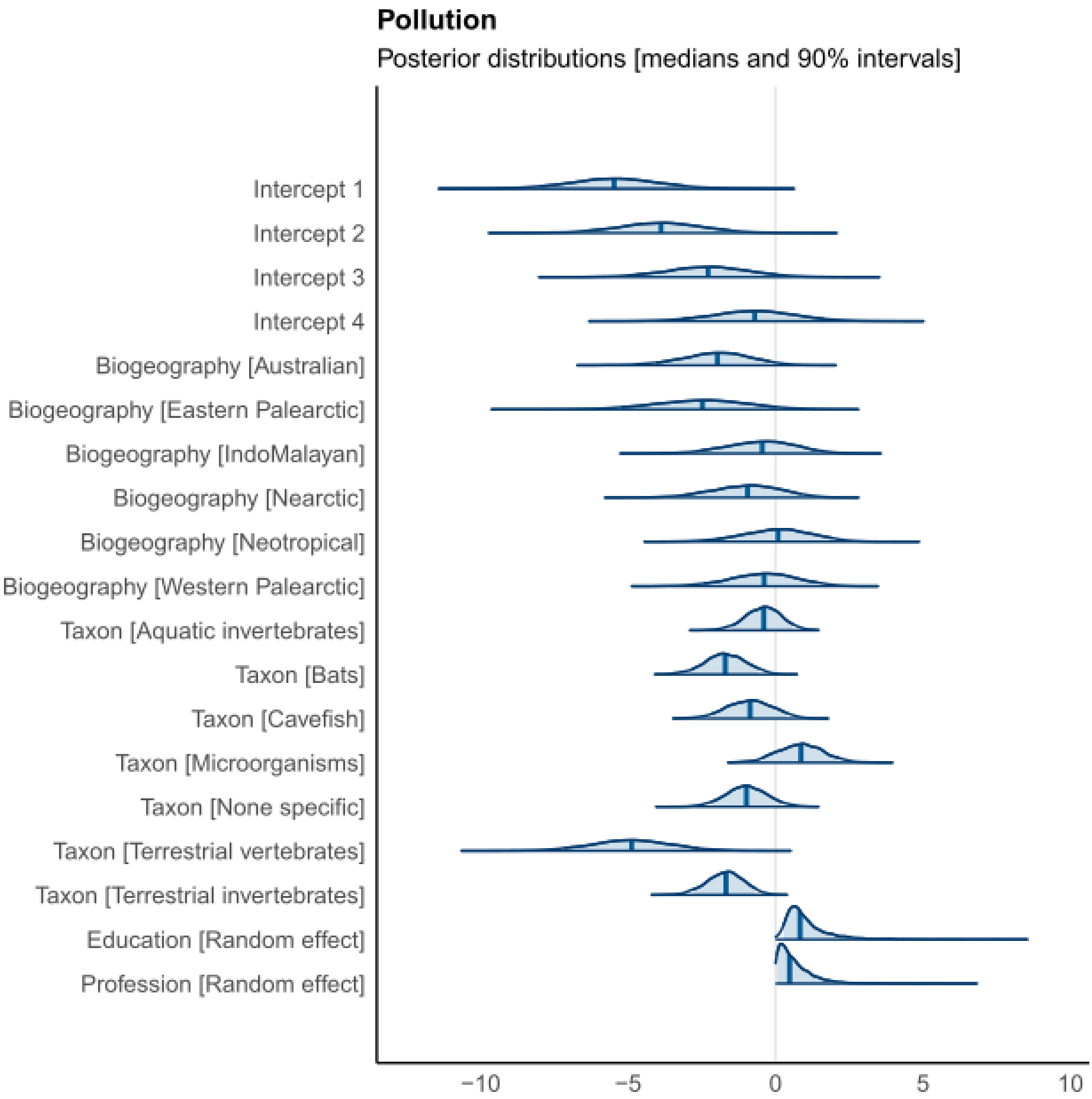

**Supplementary Figure S11**. Results of the Bayesian ordinal regression for ‘Mining’. Bayesian model density plots computed from posterior draws, with uncertainty intervals shown as shaded areas under the curves. Probability mass corresponds to 90%.

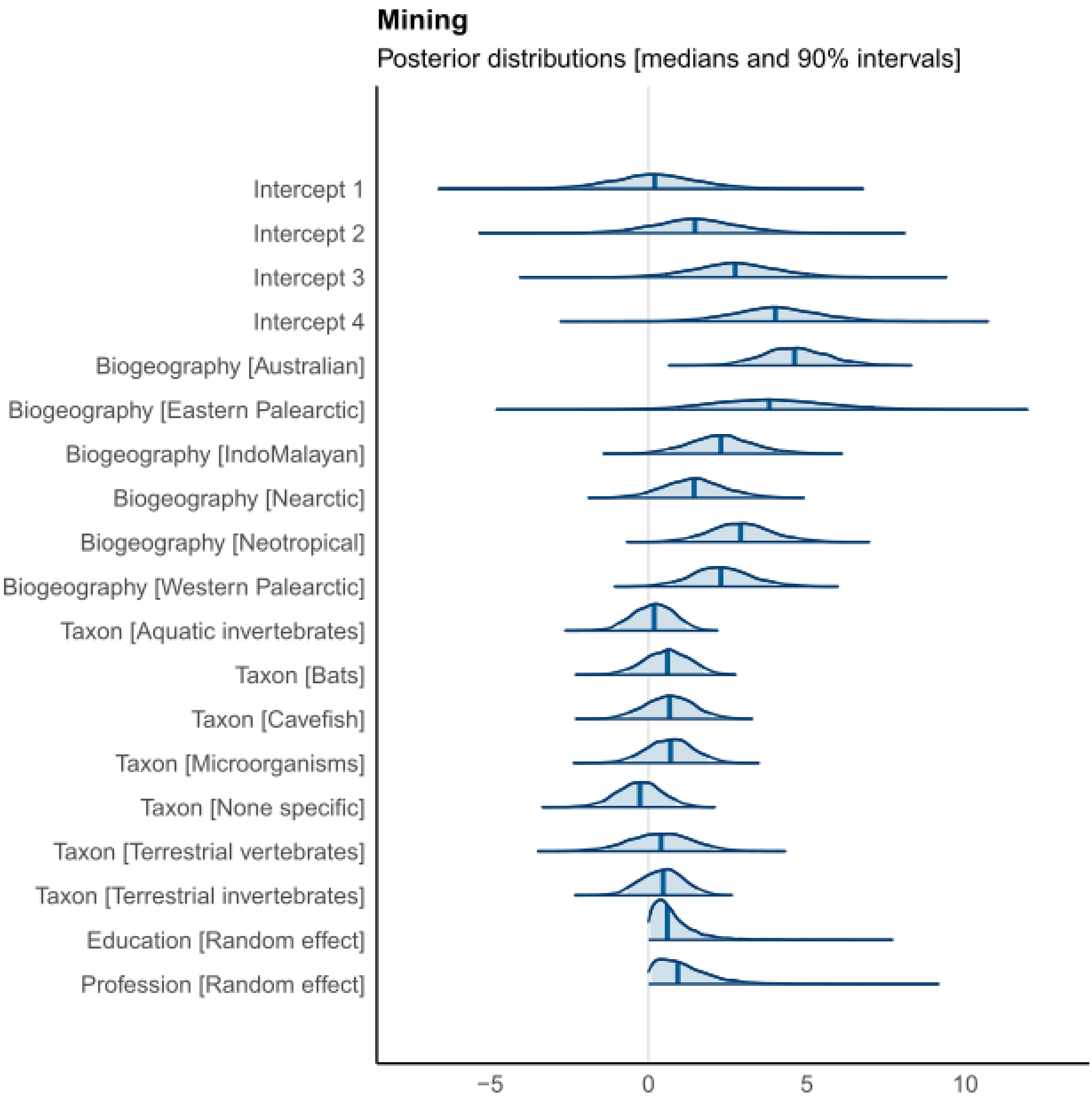

**Supplementary Figure S12**. Results of the Bayesian ordinal regression for 'Invasive species'. Bayesian model density plots computed from posterior draws, with uncertainty intervals shown as shaded areas under the curves. Probability mass corresponds to 90%.

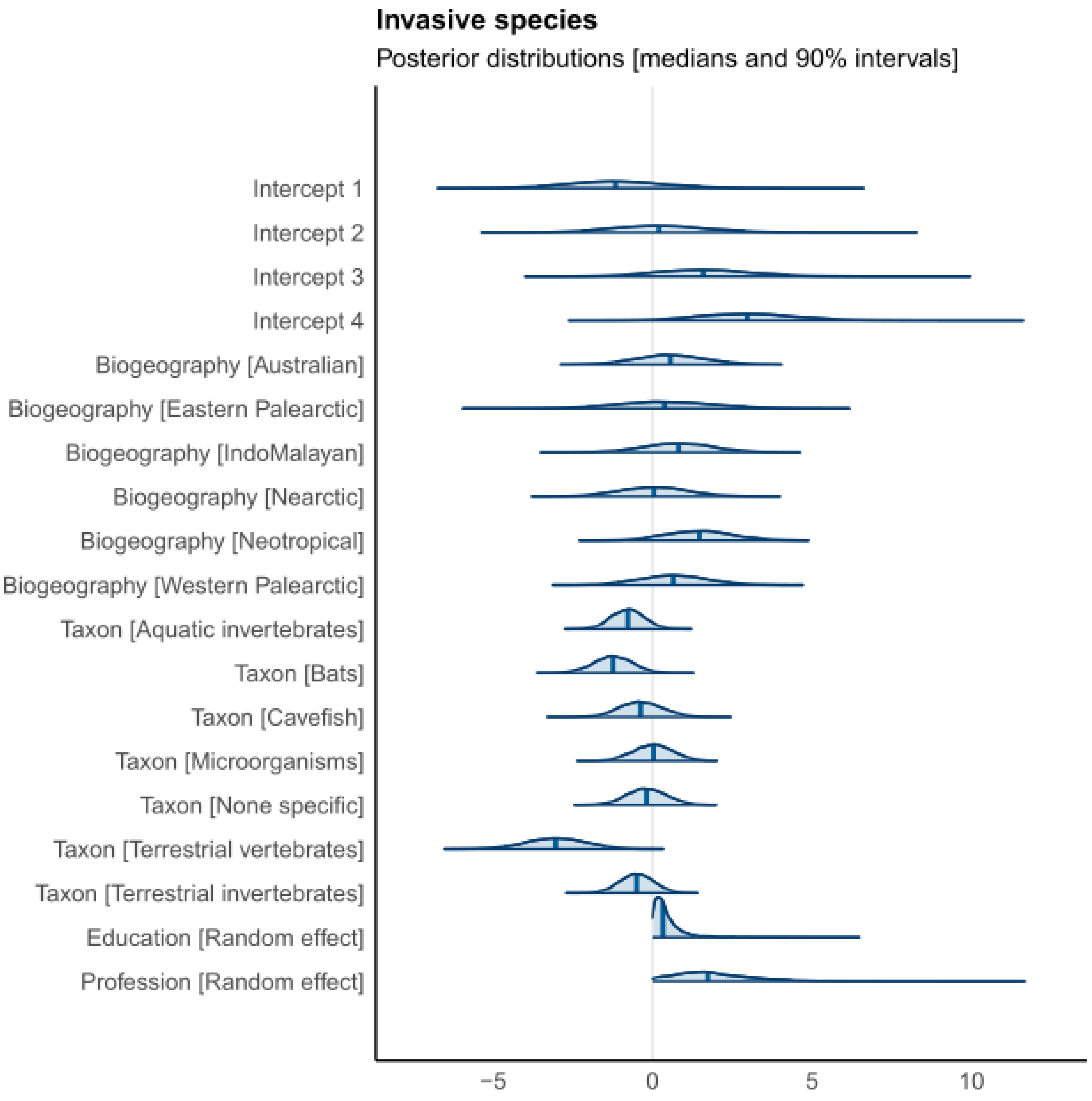

**Supplementary Figure S13**. Results of the Bayesian ordinal regression for ‘Diseases’. Bayesian model density plots computed from posterior draws, with uncertainty intervals shown as shaded areas under the curves. Probability mass corresponds to 90%.

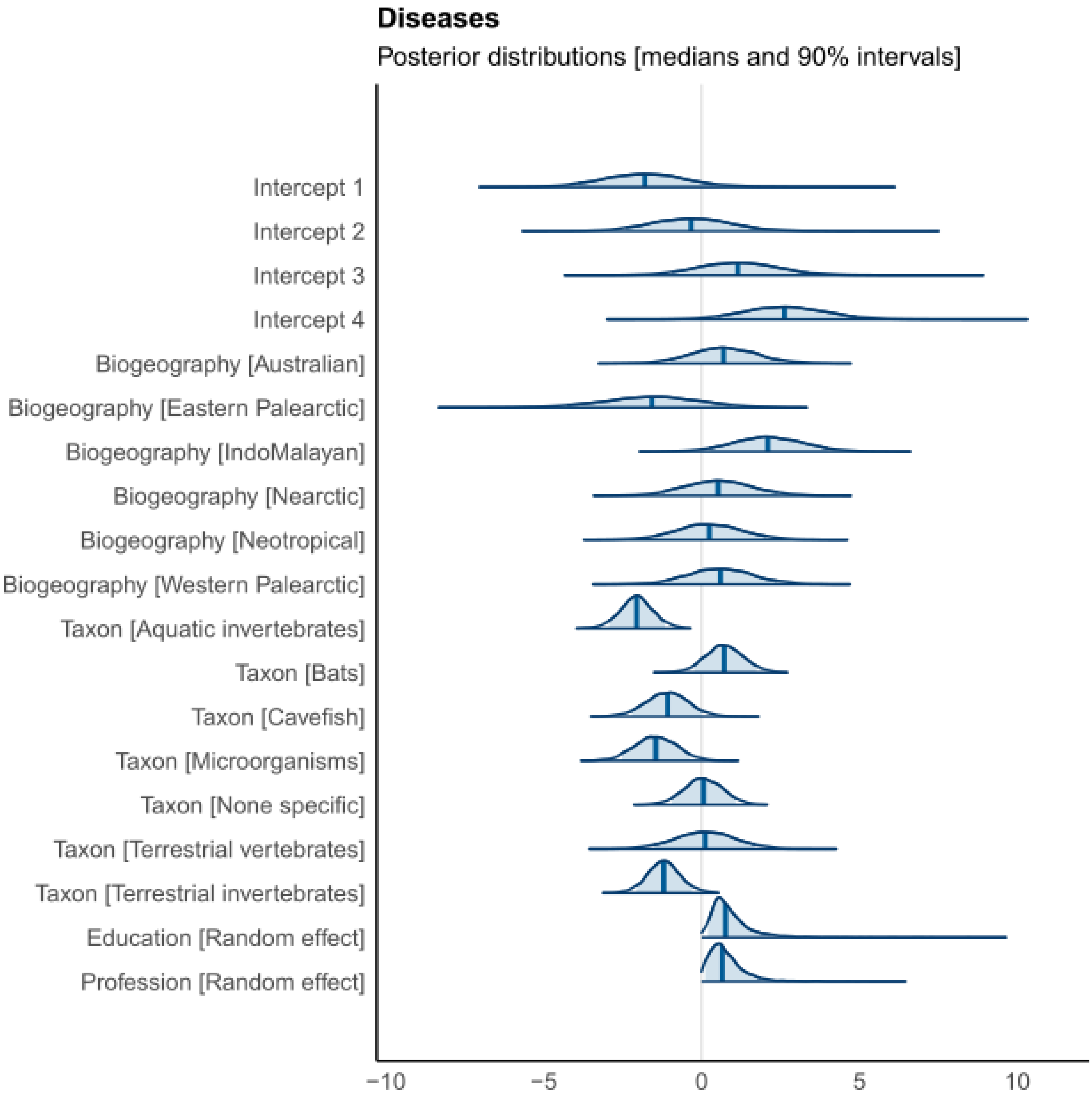

**Supplementary Figure S14**. Results of the Bayesian ordinal regression for ‘Poaching & Hunting’. Bayesian model density plots computed from posterior draws, with uncertainty intervals shown as shaded areas under the curves. Probability mass corresponds to 90%.

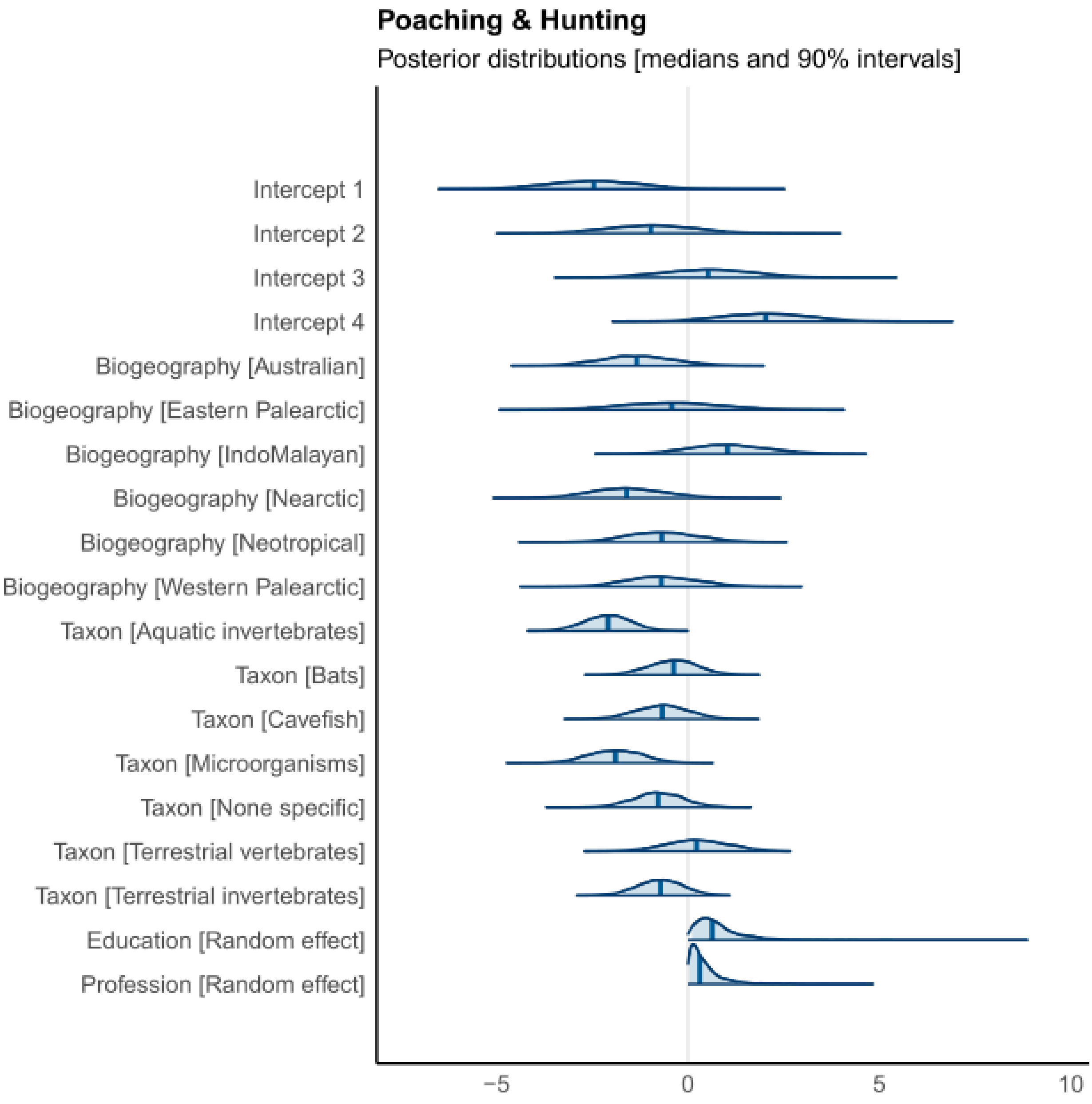

**Supplementary Figure S15**. Results of the Bayesian ordinal regression for ‘Other threats’. Bayesian model density plots computed from posterior draws, with uncertainty intervals shown as shaded areas under the curves. Probability mass corresponds to 90%.

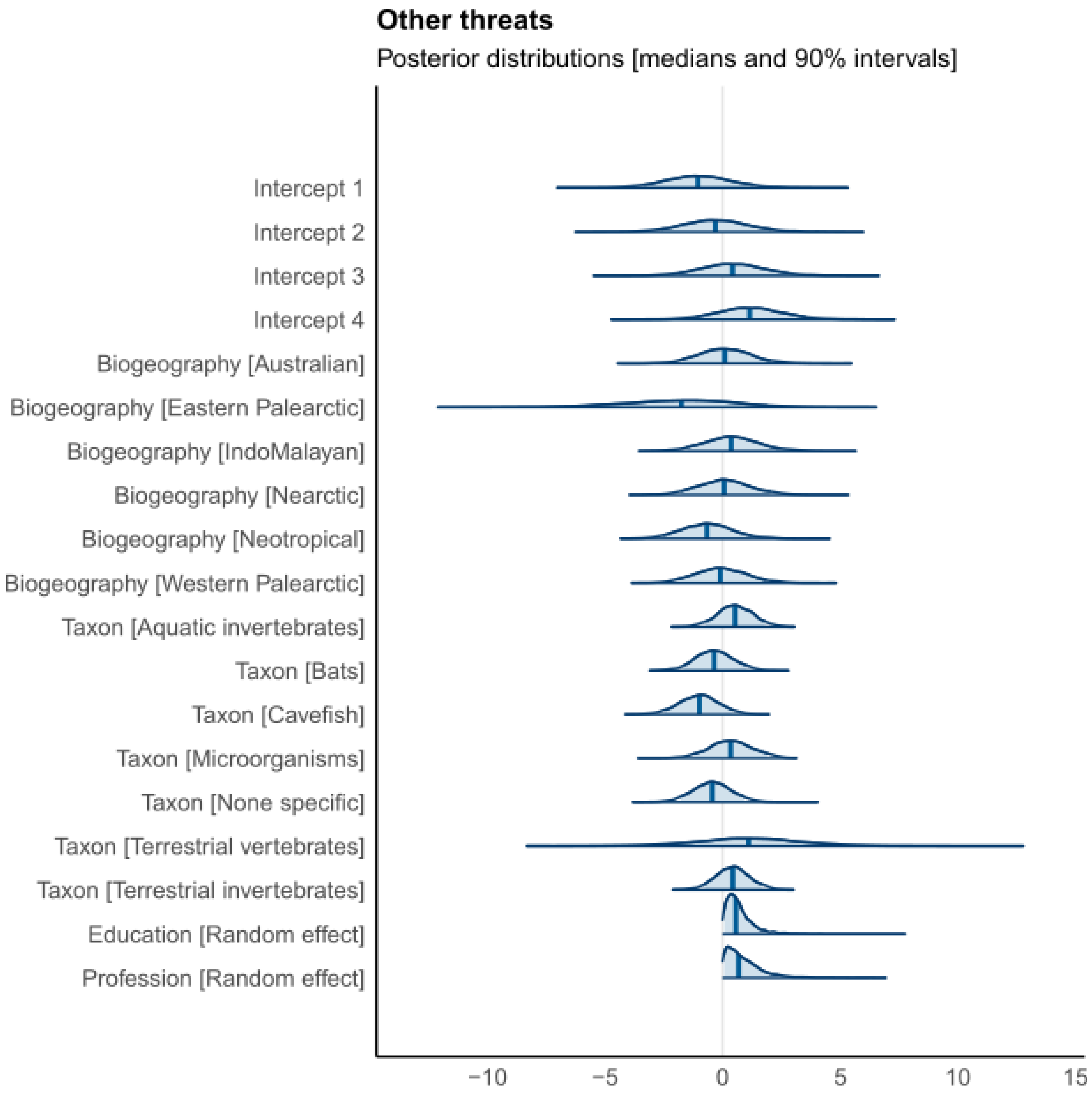

**Supplementary Figure S16**. Results of the Bayesian ordinal regression for ‘Land protection’. Bayesian model density plots computed from posterior draws, with uncertainty intervals shown as shaded areas under the curves. Probability mass corresponds to 90%.

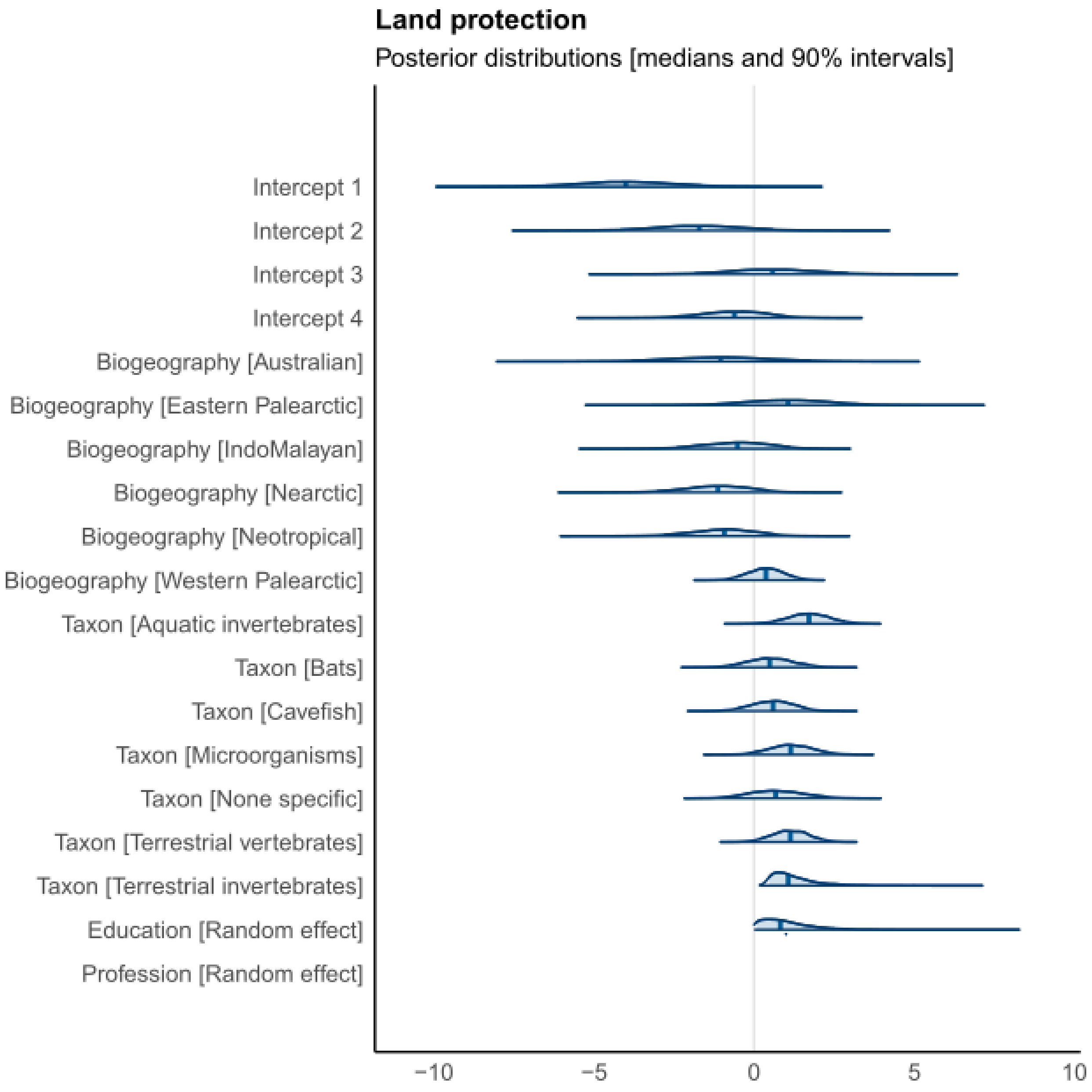

**Supplementary Figure S17**. Results of the Bayesian ordinal regression for ‘Land management’. Bayesian model density plots computed from posterior draws, with uncertainty intervals shown as shaded areas under the curves. Probability mass corresponds to 90%.

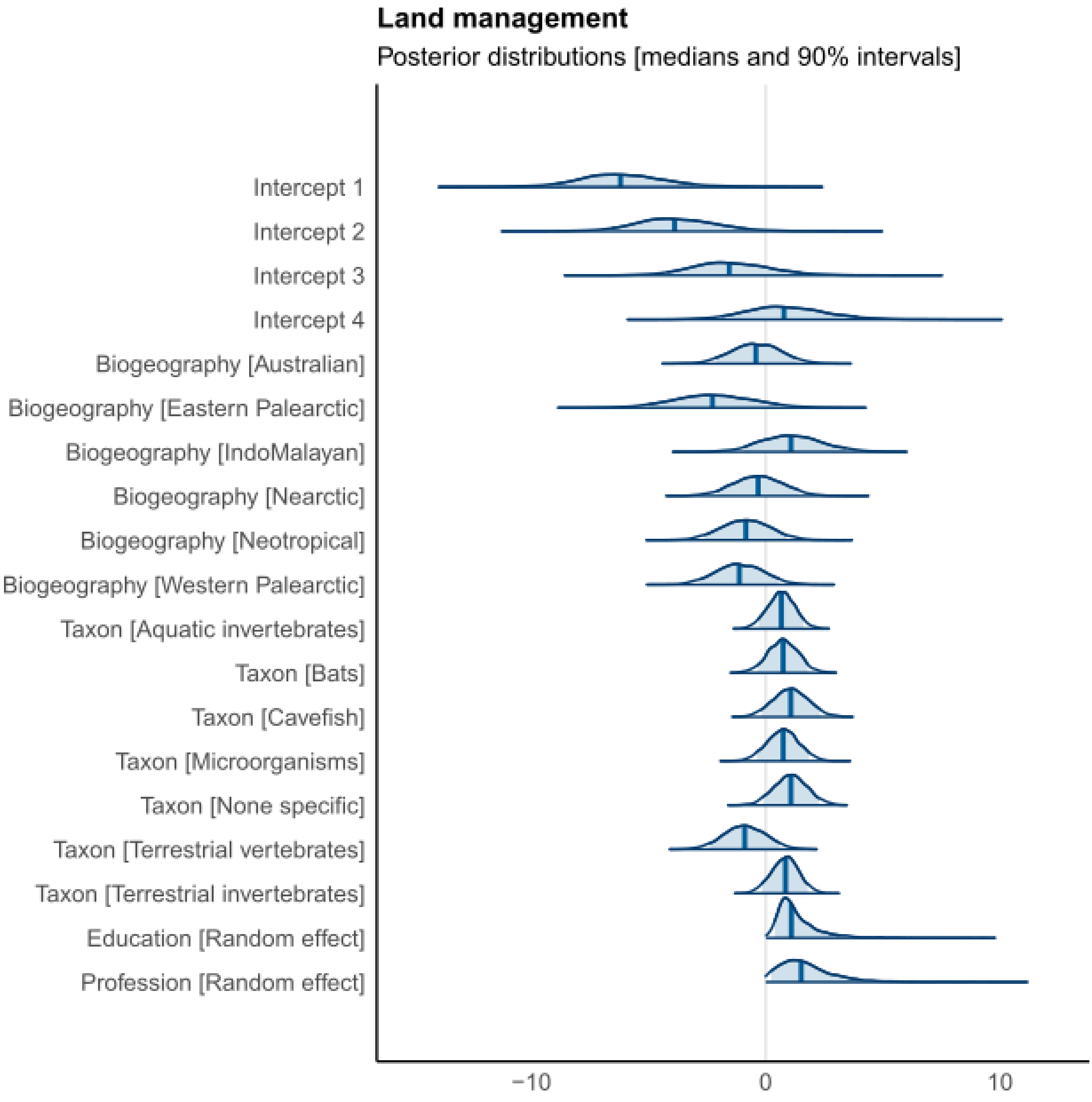

**Supplementary Figure S18**. Results of the Bayesian ordinal regression for ‘Species management’. Bayesian model density plots computed from posterior draws, with uncertainty intervals shown as shaded areas under the curves. Probability mass corresponds to 90%.

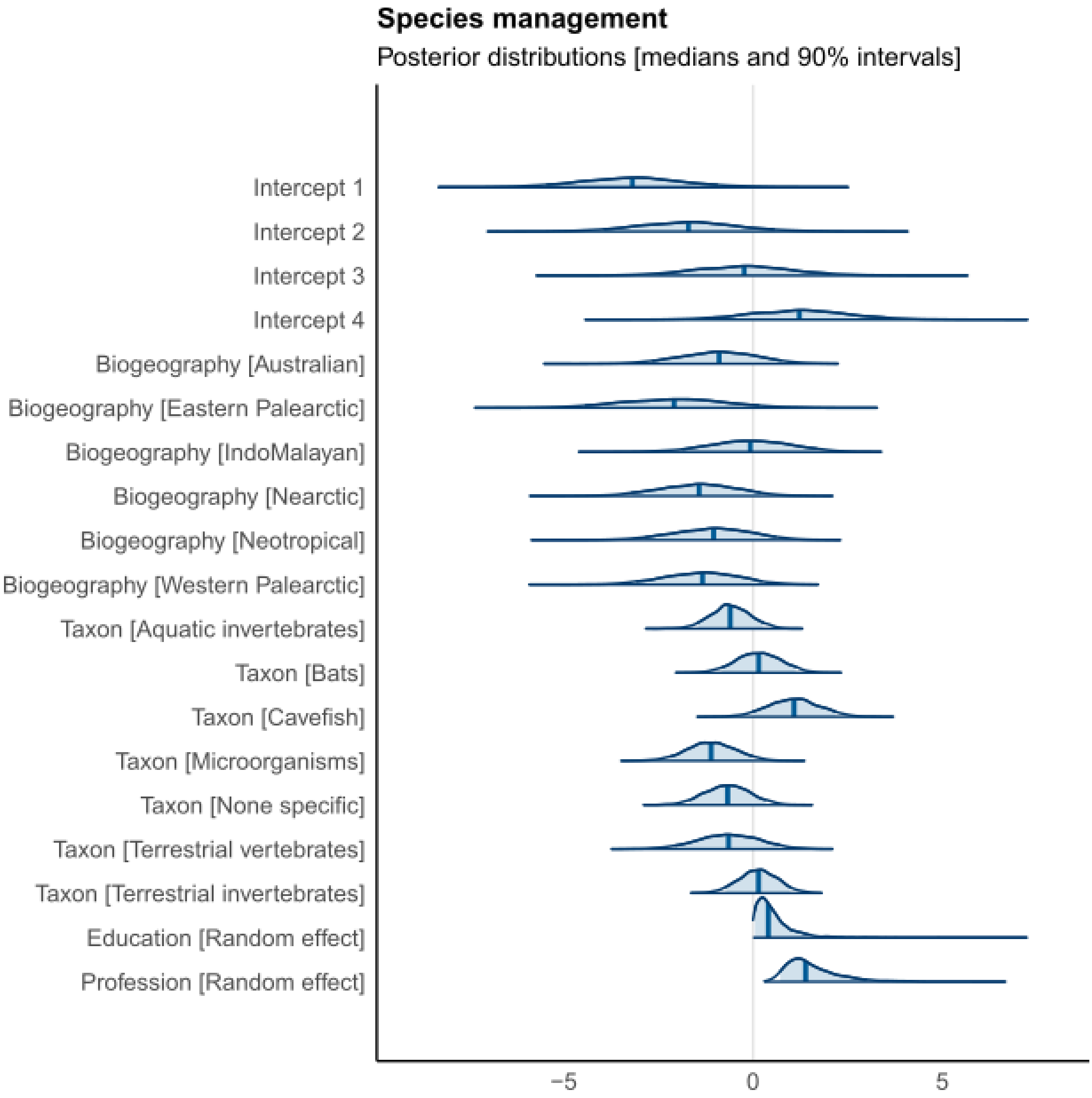

**Supplementary Figure S19**. Results of the Bayesian ordinal regression for ‘Legislation & Policy’. Bayesian model density plots computed from posterior draws, with uncertainty intervals shown as shaded areas under the curves. Probability mass corresponds to 90%.

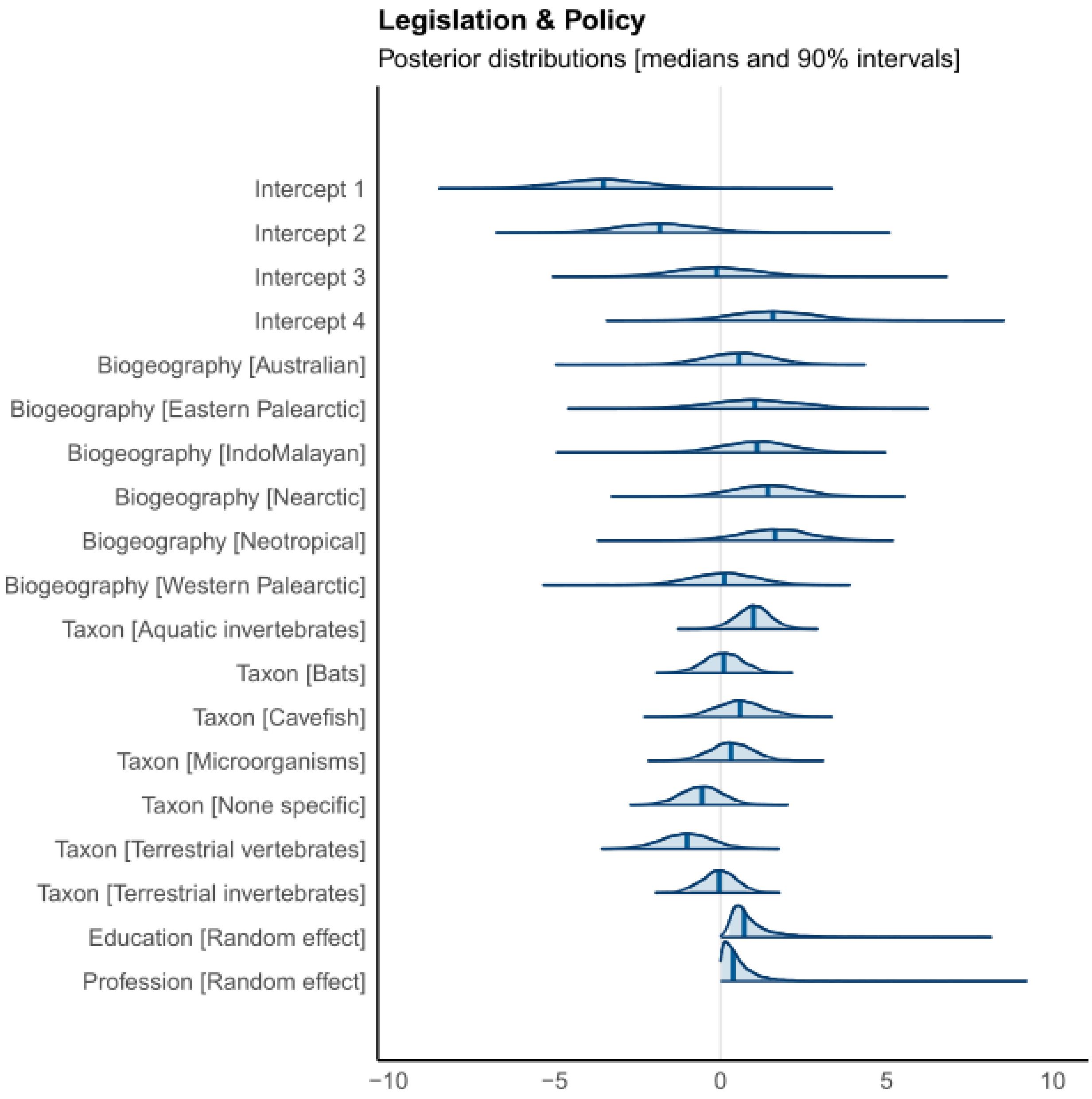

**Supplementary Figure S20**. Results of the Bayesian ordinal regression for ‘Incentives’. Bayesian model density plots computed from posterior draws, with uncertainty intervals shown as shaded areas under the curves. Probability mass corresponds to 90%.

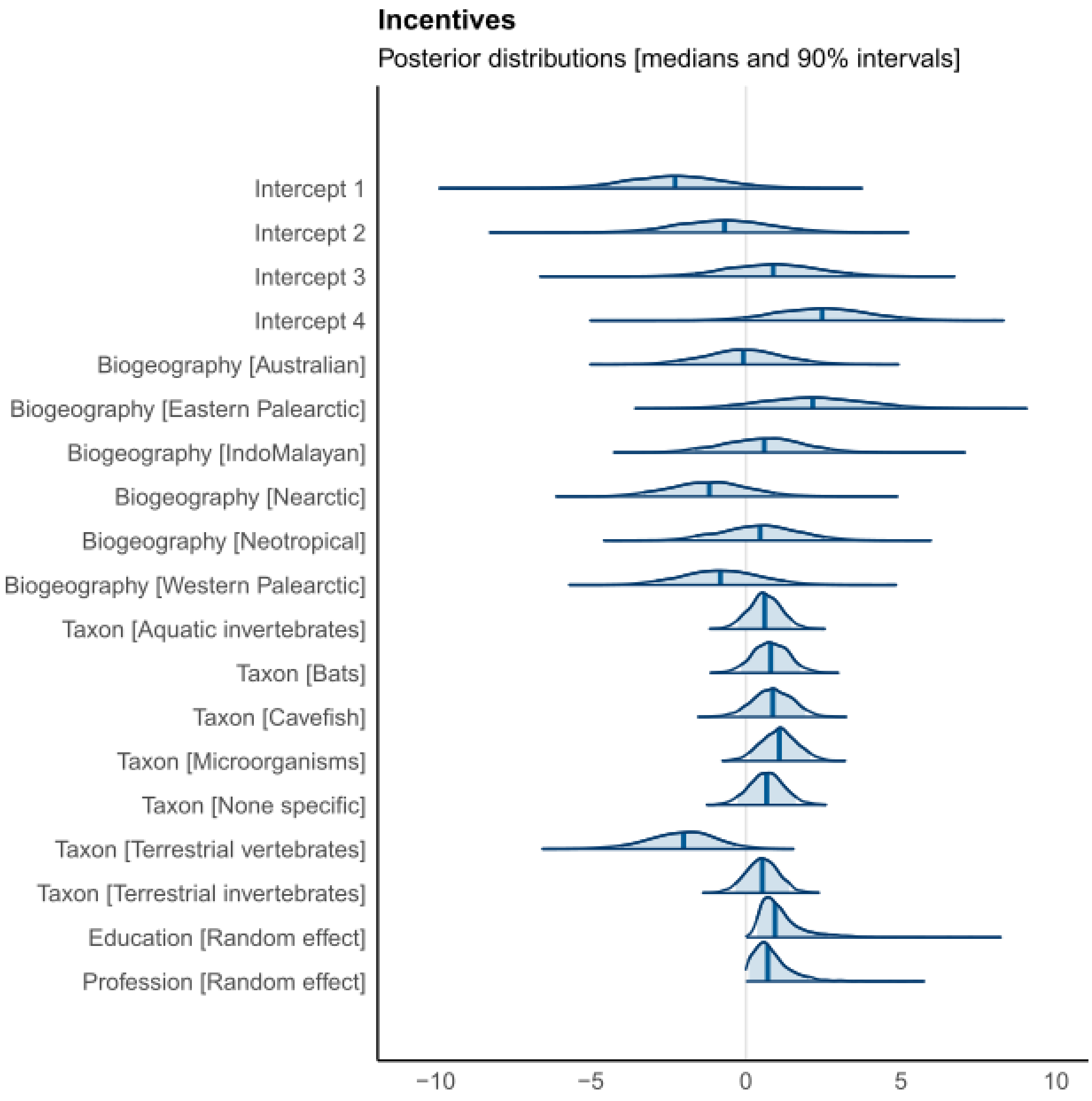

**Supplementary Figure S21**. Results of the Bayesian ordinal regression for ‘Educational activities’. Bayesian model density plots computed from posterior draws, with uncertainty intervals shown as shaded areas under the curves. Probability mass corresponds to 90%.

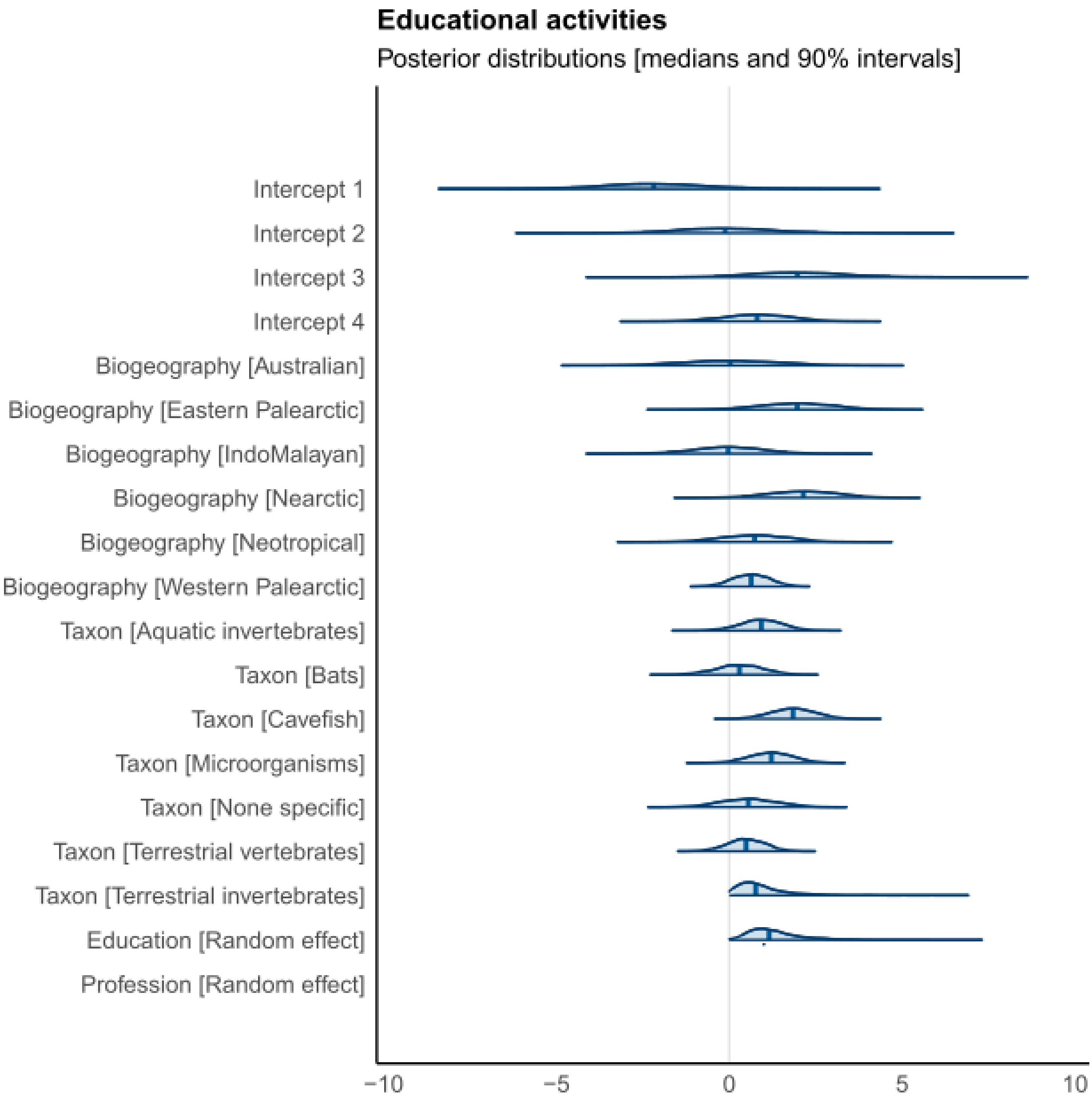
